# Phenotypic and spatial heterogeneity of brain myeloid cells after stroke is associated with cell ontogeny, tissue damage, and brain connectivity

**DOI:** 10.1101/2023.03.17.532588

**Authors:** Anirudh Patir, Jack Barrington, Stefan Szymkowiak, Dana Straus, Alessio Alfieri, Lucas Lefevre, Neil C Henderson, Karen Horsburgh, Prakash Ramachandran, Barry W McColl

**Author notes:** Authors contributed equally (Anirudh Patir, Jack Barrington).

## Abstract

Acute stroke causes substantial mortality and morbidity and provokes extensive changes to myeloid immune cell populations in the brain that may be targets for limiting brain damage and enhancing brain repair. The most effective immunomodulatory approaches will require precise manipulation of discrete myeloid cell phenotypes in time and space to avoid harmful effects of indiscriminate neuroimmune perturbation. We sought to define how stroke alters the composition and phenotypes of mononuclear myeloid cells with particular attention to how cell ontogeny and spatial organisation combine to expand myeloid cell diversity across the brain after stroke. Multiple reactive microglial states and dual monocyte-derived populations contributed to an extensive repertoire of myeloid cells in post-stroke brain. We identified important overlap and distinctions among different cell types and states that involved ontogeny- and spatial-related properties. Notably, brain connectivity with infarcted tissue underpinned the pattern of local and remote altered cell accumulation and reactivity. Our discoveries suggest a global but anatomically-governed brain myeloid cell response to stroke that comprises diverse phenotypes arising through intrinsic cell ontogeny factors interacting with exposure to spatially-organised brain damage and neuroaxonal cues.

## Introduction

Acute ischaemic stroke is a leading cause of death and disability worldwide, accounting for the greatest proportion of disease burden for all neurological disorders^1^. Recent advances in acute stroke care have improved survival rates but the majority of stroke survivors are left with long-term disability^2^, resulting in increasing numbers of people living with the disabling effects of stroke. Understanding how the brain responds to initial stroke damage to contain the injury, promote brain repair, and enhance plasticity is crucial to inform design of recovery-enhancing interventions^3,4^. Inflammatory and immune mechanisms are increasingly recognised to connect tissue damage and repair transitions throughout the body^5^, including in the CNS^6-8^, and thus are potential targets for treatments.

We and others have described how clinical and experimental stroke triggers alterations in trafficking, accumulation and phenotype of immune cells local to the stroke-injured brain tissue, and in systemic compartments^8-10^. Innate immune cells, notably macrophages of resident microglial origin as well as those derived from infiltrating blood-derived monocytes, accumulate in and around damaged brain tissue^11-14^, likely responding to an initial wave of tissue damage/distress-associated signals^15^. While some microglial/macrophage activities have potential to exacerbate tissue damage^16,17^, recent studies increasingly implicate these cells in brain protection and repair during subacute phases of stroke^18-22^ and other CNS injuries^23-25^.

Heterogeneity of mononuclear myeloid lineage cells such as microglia, monocytes, and monocyte-derived cells (MdC) can relate to their differing ontogenies, intra-population differentiation and cell plasticity, and give rise to diverse functional roles^26^. Previous studies used techniques such as transgene-encoded cell reporter mice, *in vivo* tracer labelling and bone marrow chimaeras to show mixed accumulation of broad myeloid cell populations derived from resident and infiltrating sources occurs after experimental stroke^11-14,27^. Ontogeny-dependent differences in cell phenotype and function, such as phagocytic capacity, have been suggested at broad population level^14,17,24,28,29^. Nonetheless, the above techniques are constrained for understanding deeper phenotypic heterogeneity within these cell subpopulations and relationships among them. Spatial organisation of mononuclear myeloid cell diversity in post-stroke brain is also poorly understood in part because most studies examining these cells have focussed on infarct and peri-infarct tissue. Recent scRNAseq-based studies have produced valuable new insight to cell diversity in stroke, however exclusively profiled the ipsilateral (peri-infarct) tissue therefore precluding understanding of important changes in more distant brain areas^10,30-32^. Remote effects of stroke on neurodegeneration, neuroplasticity, and glial reactivity, often in brain areas connected by white matter tracts to the primary area of damage, are likely critical to global brain network changes after stroke and consequences on motor and cognitive outcomes^33-39^.

In this study, we used complementary scRNAseq and multiplexed single molecule fluorescence *in situ* hybridisation (smFISH) approaches in an experimental stroke model (**Figure 1**). Stroke was induced by permanent middle cerebral artery occlusion (MCAO), a model not previously studied at single cell resolution, and important given that half of stroke patients do not achieve adequate tissue reperfusion even with recanalisation^40^. We focussed analysis at 3 d post-MCAO, a critical juncture in the subacute phase characterised by emergence of adaptive mechanisms responding to initial damage, and studied cellular changes in brain areas local and distant to the focal cortical injury. We present multiple novel findings showing that experimental stroke substantially expands the diversity of mononuclear myeloid cells through spatially-dependent combinations of altered cell ontogeny, differentiation, and reactivity. We identify location-dependent phenotypes such as *Gpnmb*-expressing MdC dominant in intra/peri-infarct areas and *Ccl* chemokine-enriched microglia in remote but connected brain areas and along the white matter connecting tracts. Multiple microglial reactive states are identified responding to stroke, some with similarities to those found in other pathologic conditions, and which are spatially heterogeneous. Some reactive microglia shared partial but substantial gene expression identity (e.g. *Apoe, Ctsb, Fabp5, Plin2, Spp1*) with non-resident stroke-induced macrophages derived from infiltrating monocytes and these were closely intermixed on a micro-anatomical scale but restricted to peri-infarct areas. Our data also reveal that proliferation selectively within microglia (occurring proximal and in distant connected regions to primary injury) and dual differentiation trajectories within Mo/MdC (accumulating only proximal to injury) contribute to the expanded mononuclear myeloid phenotypic diversity after stroke. Our findings bring new insight, increased clarity on existing uncertainties, and prompt new questions around the global brain cellular response to stroke from the perspective of microglia, macrophages, and related myeloid cell types.

**Figure 1.**
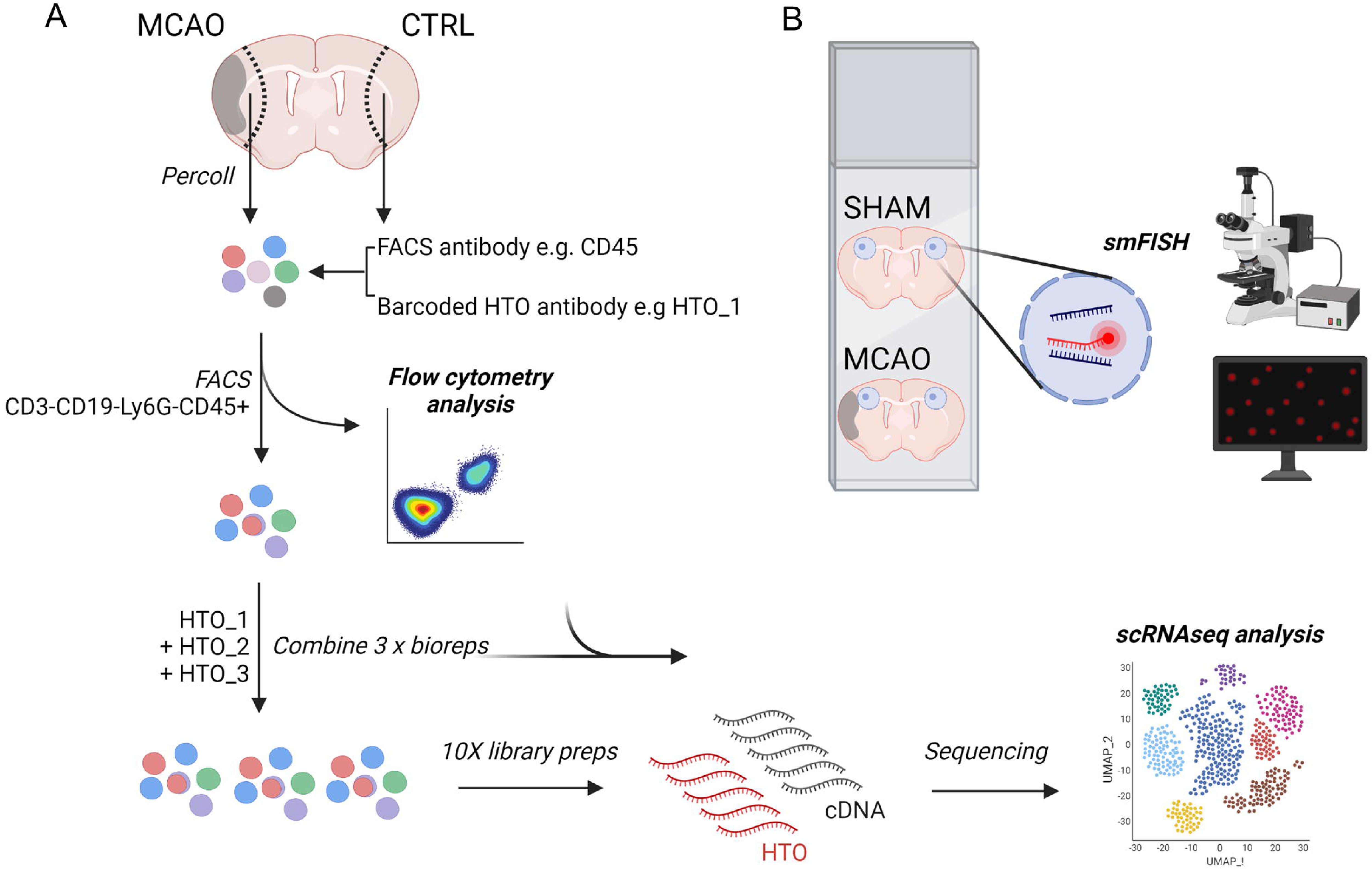
Experimental workflow. (A) Brain cell suspensions were prepared from ipsilateral (MCAO) and contralateral (CTRL) hemispheres by density-gradient centrifugation and cells were labelled with FACS antibodies to surface markers and with hash-tag oligonucleotide (HTO)-conjugated antibodies to barcode individual samples and enable sample multiplexing. Cells were sorted to exclude the majority of lymphocytes and granulocytes and concurrent cytometry data acquired for analysis. Sorted bioreplicate samples were pooled for each experimental condition (MCAO or CTRL) and single cell cDNA libraries prepared for sequencing. Data were processed and analysed using a Seurat workflow with additional tailored methods as described in Methods. (B) Brain sections from sham-operated and MCAO-operated mice were labelled with fluorescence oligonucleotide probes and quantitative image analysis conducted as described in Methods. smFISH, single molecule fluorescence *in situ* hybridisation.

## Results

### Flow cytometric overview of MCAO-induced immune cell changes

The scRNAseq experimental workflow comprised sorting myeloid-enriched brain cell suspensions 3 d after MCAO from multiple donor mice identified by “cell hashing” for droplet-based single-cell RNA sequencing and computational analysis, summarised in **Figure 1A**. We acquired flow cytometric data during the cell sorting procedure (see below) to provide a contextual overview of changes to absolute abundance of major immune cell subclasses in response to MCAO. A largely homogeneous CD45^+^CD11b^+^ population consisting almost entirely of CD45^lo^ cells was observed in brain cell suspensions from the hemisphere contralateral to MCAO (hereafter referred to as the CTRL condition) (**Figure 2A**). Two additional CD45^hi^ populations were evident in ipsilateral hemisphere samples (hereafter referred to as the MCAO condition), comprising CD11b^+^Ly6G^+^ (neutrophils) and CD11b^+^Ly6G^-^ cells. We and others have shown previously that CD45 surface expression levels can adequately distinguish resident microglia from non-parenchymal myeloid cells under certain circumstances^41^. Therefore, we considered CD45^lo^CD11b^+^Ly6G^-^ cells as resident microglia and the CD45^hi^CD11b^+^Ly6G^-^ population as blood-derived monocytes (Mo) and monocyte-derived cells (MdC) recruited in response to MCAO (full gating scheme for quantification is shown in **Figure S1**). MCAO caused a significant increase in the number of total CD45^+^ cells, total CD11b^+^ cells, Ly6G^+^ granulocytes, microglia, and CD11b^+^CD45^hi^Ly6G^-^ Mo/MdC, indicating marked innate immune cell infiltration and an overall increased myeloid cell abundance (**Figure 2B**). In contrast, there were negligible CD3^+^ and CD19^+^ cells in either condition indicating limited parenchymal T cell and B cell accumulation (**Figure 2A, C**). The proportions of the cell subsets (microglia, Mo/MdC and neutrophils) comprising the CD11b^+^ parent population changed significantly after MCAO, showing that the composition within the myeloid cell compartment was altered by MCAO (**Figure 2D**). The median CD45 intensity of cells specifically within the microglial population, a measure of their broad reactive amplitude, was significantly greater (∼45%) in MCAO samples although remained ∼10-fold lower than that of the clearly separated CD45^hi^ infiltrating myeloid population (**Figure 2E, F**), as we found previously^41^. There was also broader dispersion of individual microglial CD45 intensities within MCAO samples (**Figure 2F**), reflected by a greater coefficient of variance (**Figure 2G**), suggesting greater heterogeneity within the overall microglial population.

**Figure 2.**
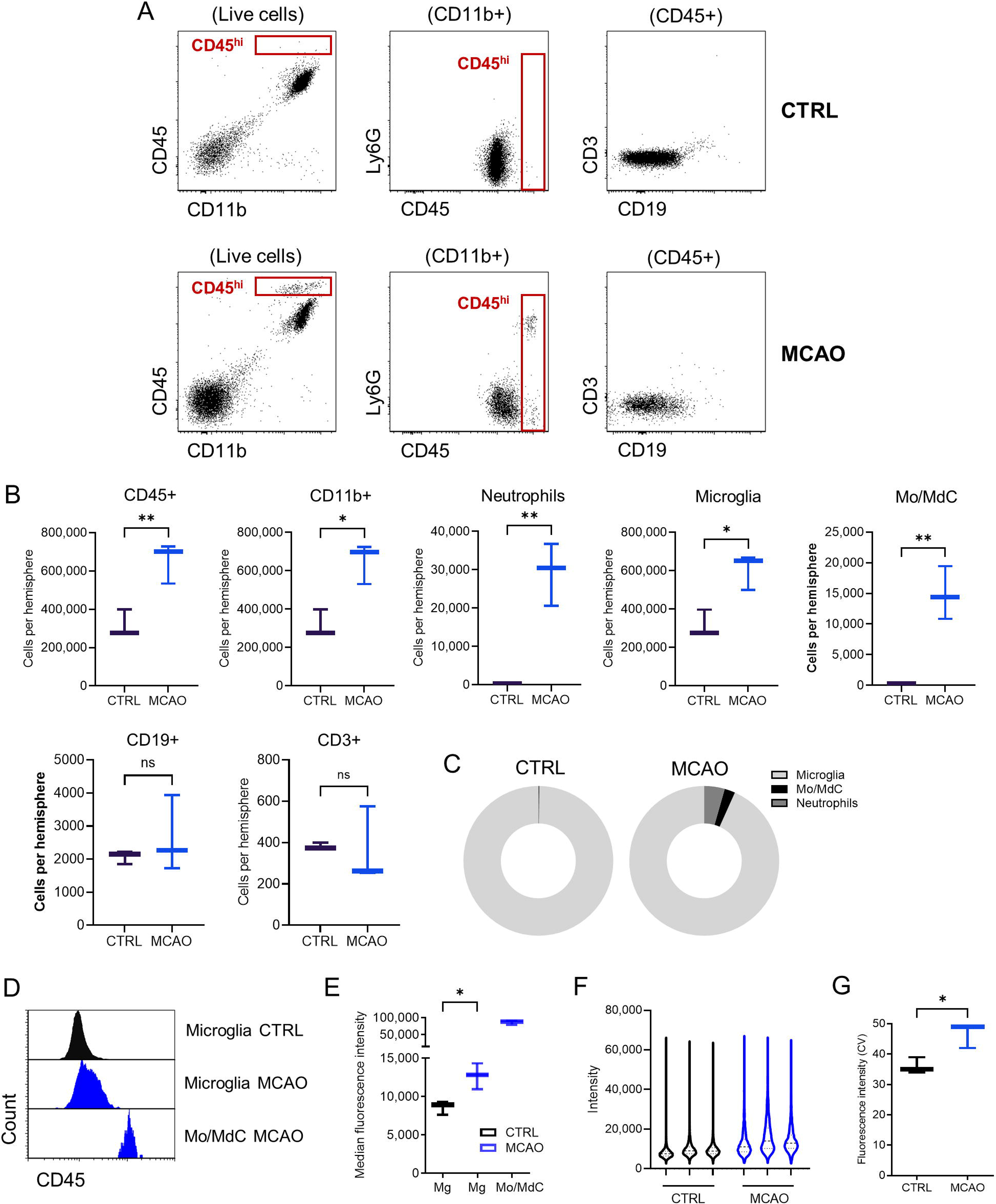
Flow cytometry analysis of myeloid cell composition. Brain cell suspensions prepared from ipsilateral (MCAO) and contralateral (CTRL) hemispheres 3 d after MCAO were analysed by flow cytometry using a gating strategy as described in **Figure S1**. (A) Representative dot plots for selected surface marker combinations showing the altered pattern of cell accumulation between MCAO and CTRL samples. Text in parentheses above dot plots indicates the pre-gated cell population. (B) Quantification of major cell populations defined according to the gating strategy as described in **Figure S1**. *P<0.05, **P<0.01 Welch’s t-test, n = 3 per group; boxplot whiskers denote min and max values. (C) Composition of the myeloid cell compartment for major cell subclasses (expressed as a proportion of CD11b^+^ cells) in MCAO and CTRL samples. (D) Representative histograms and (E) quantification of CD45 intensity within pre-gated cell populations. *P<0.05 Welch’s t-test, n = 3 per group; boxplot whiskers denote min and max values. (F) Violin plots showing the dispersion of CD45 intensity values across all cells within the pre-gated microglial population for each sample. (G) Comparison of the coefficient of variance (CV) for the microglial CD45 intensity distributions in (F). *P<0.05, Welch’s t-test, n = 3 per group. Mo, monocytes; MdC, monocyte-derived cells.

### Cell sorting and hashtag oligonucleotide (HTO) labelling validation

To enrich brain cell suspensions prepared from cortical samples ipsilateral and contralateral to MCAO for myeloid mononuclear cells, we sorted on cells negative for pan-lymphocyte (CD9 and CD19) and granulocyte (Ly6G) cell surface markers and positive for CD45 (**Figure S2A**). Cell suspensions for sorting were co-incubated with hashtag oligonucleotide (HTO)-conjugated antibodies (that recognise the CD45 antigen ubiquitous on immune cells), enabling demultiplexing of samples pooled from different donor mice and multiplet identification^42^. We pre-validated that the HTO-conjugated antibody and the fluorochrome-conjugated CD45 antibody used in the cocktail for cell gating and sorting would not out-compete each other by confirming co-staining of microglia and bone marrow immune cells with two CD45 antibodies (of the same clone as HTO and sort cocktail antibodies) each identifiable by distinct fluorochromes. Labelling of cells with both CD45 antibodies at a similar intensity was evident across and within samples (**Figure S2B, C**), supporting the use of HTO-conjugated antibodies to demultiplex samples and, in future, to use HTO-derived sequencing output as a proxy to infer quantitative expression levels of cell surface cognate antigen (i.e. CITE-SEQ)^43^.

### Altered mononuclear myeloid cell composition 3 days after MCAO

Using the aforementioned procedure, mononuclear myeloid cells were isolated by FACS and mRNA libraries were prepared, sequenced and aligned to generate gene expression matrices. Low quality cells were filtered out (based on the number of features expressed, library size and mitochondrial content). We identified a small proportion (20%) of multiplets using the HTO barcodes and removed these prior to analysis (**Figure S3A-C**). Clustering cells according to normalised HTO expression after multiplet removal organised cells according to their donor of origin (**Figure S3D, E**). Donor variation was accounted for using canonical correlation analysis (CCA), which facilitates the integration of donor-derived data based on similar cell types. The resultant integrated dataset comprised of 5,213 cells.

To cluster cells while capturing the heterogeneity or variation within the data, genes were first reduced to the 47 most significant (*P* < 0.05) principal components (PC) using principal component analysis. Thereafter, an unbiased graph-based algorithm was used to cluster cells sharing similar PC profiles (a proxy of their transcriptome). The analysis identified 22 cell clusters organised into four major components when projected onto a two-dimensional space using uniform manifold approximation and projection (UMAP) ^44^ (**Figure 3A**). Cells were also labelled according to their experimental condition, either originating from the hemisphere ipsilateral (MCAO) or contralateral (CTRL) to arterial occlusion (**Figure 3B**), and their specific mouse donor determined from the HTO barcodes (**Figure 3C).** Overlaying mouse donors with expression clusters showed that each of the 22 clusters comprised cells from each donor mouse (**Figure 3C, D),** thus demonstrating replication of scRNAseq expression profiles across multiple animals. We annotated the 22 cell clusters by surveying the expression of canonical marker genes for various immune cell classes combined with the expression of the most variable genes for each cluster (**Figure 3D-E, Figure S4, Table S1**). As anticipated from the enrichment for mononuclear myeloid cells during cell sorting, almost all clusters were of myeloid origin and there was no identifiable granulocyte cluster. We detected a cluster of NK cells (*Klrb1c*^+^), which would not be expected to be gated out using our antibody cocktail, and small clusters of B cells (*Ighm*^+^) and T cells (*Trac^+^*), likely present because small proportions of these cells express low levels of the CD19 and CD3 surface antigens. All remaining 19 clusters were clearly identifiable as myeloid based on the expression of mononuclear myeloid lineage-defining genes such as *Csf1r* combined with broad subpopulation-selective myeloid genes (e.g. *P2ry12*, *Ccr2, Flt3*). Broadly, these clusters were identified as microglia, perivascular/border-associated macrophages (PVM/BAM), dendritic cells (DC), monocytes, and monocyte-derived cells (**Figure 3D**, **Table S2**).

**Figure 3.**
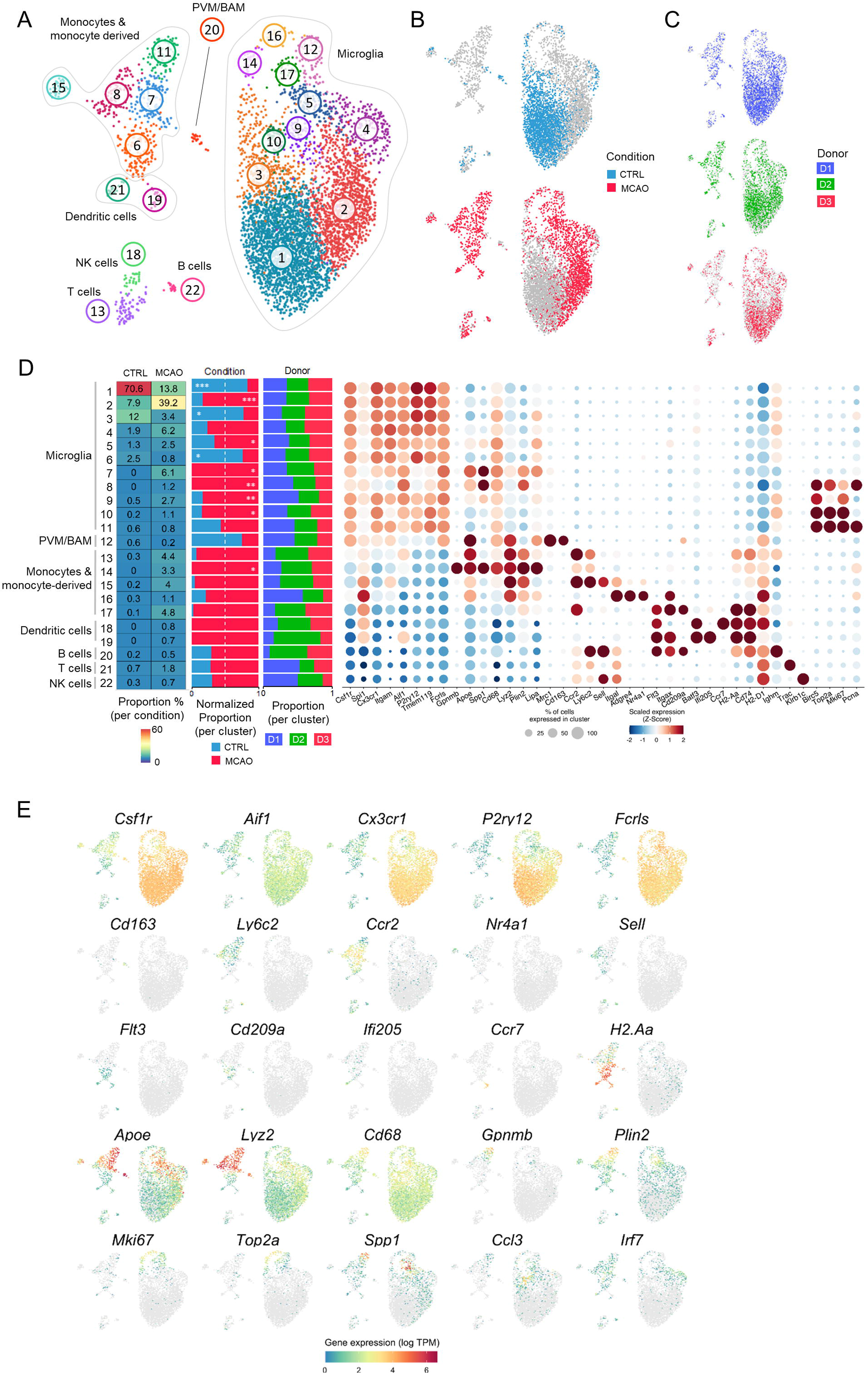
Cell clustering and annotation. (A) After quality control and processing of scRNAseq data, 5,213 cells were clustered into 22 clusters based on their transcriptomic profile and projected in two dimensions using UMAP to highlight visually their similarities and differences. (B) UMAPs show cells derived from the CTRL (top) and MCAO (bottom) experimental conditions and (C) cells derived from each of the three donor mice. (D) From left to right the panels display for each cell cluster (rows) their proportion within each condition, their distribution across condition, their distribution across donors and the expression of selected markers shown as a dot plot. For the dot plot, the colours represent the mean expression value which is scaled across clusters and the size of the dot indicates the number of cells of the cluster (column) expressing the gene (row). Annotation of major cell groupings in (A) and (D) is based on annotation of individual clusters (see **Table S2**) (E) Expression profile of selected immune cell marker genes projected by UMAP. *P< 0.05, **P <0.01, ***P<0.001. PVM, perivascular macrophage; BAM, border macrophage.

A large group of brain-resident microglia, comprising multiple sub-clusters (described below) was evident by a high expression of signature genes including *P2ry12, Tmem119, Gpr34, and Fcrls*. These cells were spatially segregated on the UMAP from other myeloid cell types with no or negligible expression of these genes. Ly6C^hi^ (cluster 8) and Ly6C^lo^ (cluster 15) monocytes were clearly distinguished by their relative expression of genes such as *Ccr2, Ly6C, Nr4a1 and Adgre4*. Clusters juxtaposed to Ly6C^hi^ monocytes in the UMAP projection implicated these as differentiated derivatives of monocytes progressing along macrophage (cluster 7, 11) and dendritic cell (cluster 6) trajectories based on both canonical gene expression (e.g. *Apoe, Lyz2* for macrophages; *Flt3, Itgax* for dendritic cells) and expression gradients indicative of their direct maturation from monocytes (e.g. declining but detectable *Ccr2* and *Sell* expression). Two clusters of dendritic cells were identifiable as conventional dendritic cells (cDC, e.g. high expression of *Batf3, Irf8, Ifi205*) and migratory DCs (MigDC, e.g. high expression of *Ccr7* and *Anxa3*). The relationships among these monocyte-derived and DC clusters are examined in further detail below. An additional cluster of *P2ry12*^lo^*Fcrls*^+^ cells distinct from microglia expressed high levels of *Mrc1, Cd163* and *Pf4,* therefore characteristic of perivascular macrophages (cluster 20). Several clusters of cells expressed high levels of cell cycle-associated genes (e.g. *Mki67, Top2a*), primarily restricted to those of microglial identity (clusters 12, 14, 16, 17).

We assessed the relative abundance of cells derived from MCAO or CTRL samples within each cluster and the representation of each cell cluster within MCAO and CTRL conditions (**Figure 3D**). Microglia, when considered collectively, comprised the vast majority (∼98%) of CTRL sample-derived cells, whereas this declined to <80% of cells in the MCAO sample mainly due to the accumulation of monocytes (∼17.5%) and DCs (∼1.5%). Cluster 1 cells were the most abundant (71%) in the CTRL condition but declined to 14% in the MCAO condition, notable because the cells in this cluster expressed the highest levels of many genes (e.g. *P2ry12, Tmem119,* and *Gpr34*), reflecting the microglial homeostatic states. When assessing the presence of MCAO- or CTRL-derived cells within each cell cluster, the most notable observation was that the composition of 13 of 19 myeloid clusters, including both microglial and non-resident monocyte/MdC/DC clusters was heavily dominated by MCAO-derived cells, therefore, indicating their emergence primarily in the MCAO sample. Some cell clusters (7, 8, 14, 18, 19) were exclusively comprised of MCAO-derived cells. Collectively, these data show a marked expansion in the diversity of brain myeloid cell ontogenies and phenotypes 3 d after experimental stroke

### Heterogeneity of reactive microglia 3 days after MCAO

We explored in greater depth the impact of MCAO on brain-resident microglial reactive heterogeneity by sub-clustering on the population of *P2ry12^+^Fcrls^+^* cells, thus excluding non-parenchymal border macrophages (*P2ry12-Fcrls^+^*) and non-resident immune cells (*P2ry12-Fcrls-*). As a result, 16 microglial cell clusters (cMG) were identified (**Figure 4A**) containing multiple groups of homeostatic and reactive states (**Figure 4B-D, Table S3**). Next, an unbiased approach, WGCNA on the 3,890 genes that were differentially expressed across microglia (**Figure S5, Table S4**), was adopted to identify gene coexpression modules (gMG) associated with these microglial clusters. These gene modules signify groups of genes with highly similar expression profile across cells. To improve the signal-to-noise ratio of scRNAseq data and create a balanced gene correlation network, pseudo-bulk samples were generated for each cluster before conducting WGCNA (see Methods). 18 gMGs were identified (**Figure 4D, Figure S6, Table S5**) and enrichment analysis was performed on each of the gMGs based on gene sets of Reactome, KEGG, and GO terms to explore biological pathway representation (**Figure 4D, Table S6)**.

**Figure 4.**
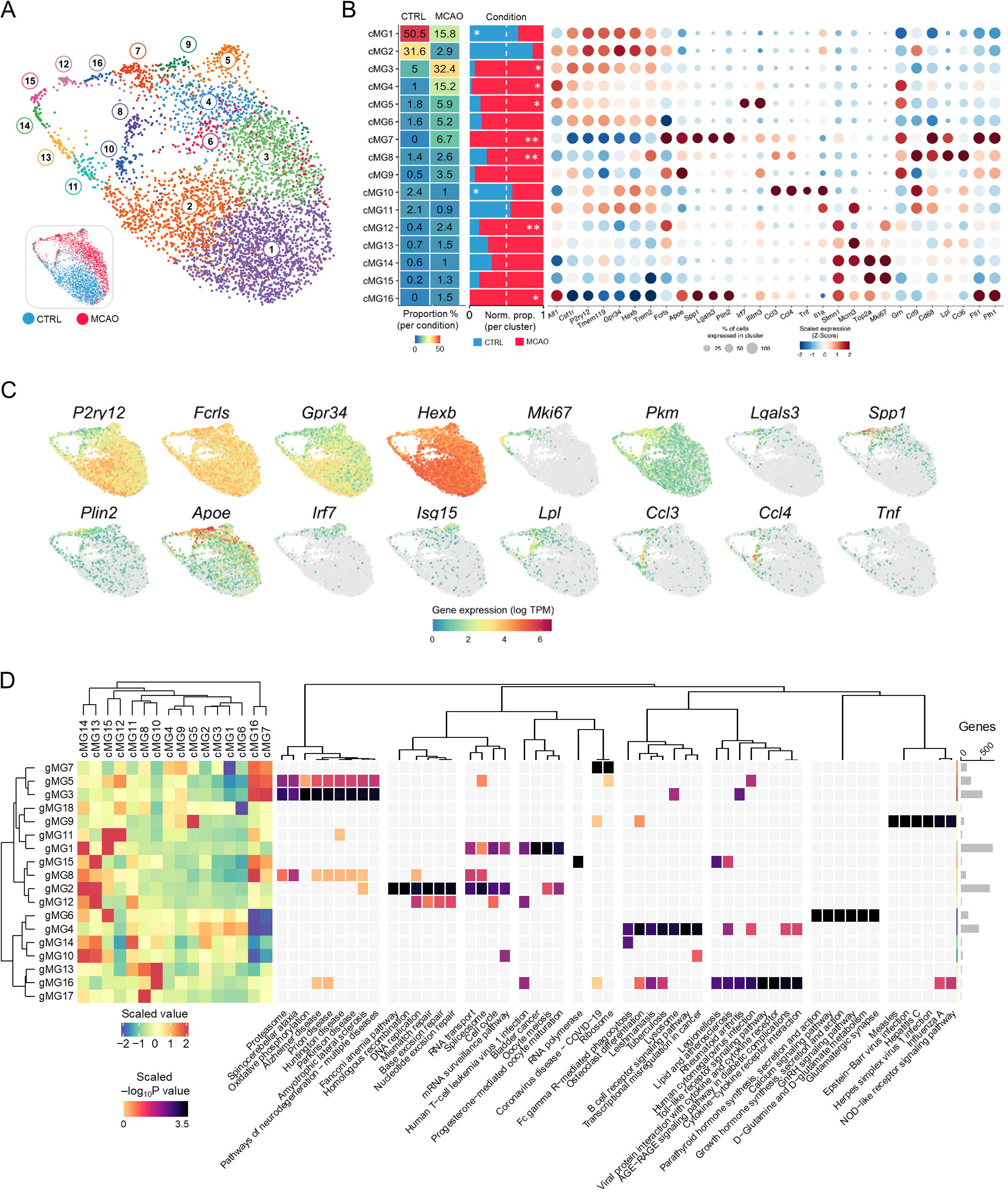
Multiple microglial reactive states associated with MCAO. (A) Microglia were sub-clustered from the entire dataset and based on their transcriptomic profile projected in 2D using UMAP. Inset UMAP shows the MCAO and CTRL conditions from which sub-clustered microglial cells were derived from. (B) From left to right, for each cell cluster (rows) panels show the proportion of all cells contributing to each of the MCAO and CTRL conditions, the proportion of cells within the cell cluster derived from each condition, and a dot plot of the expression of marker genes. For the dot plot, colours represent mean expression value scaled across clusters and point size indicates number of cells expressing the gene (column) within each cluster (row). (C) Expression of selected marker genes projected on the cell UMAP. (D) Using gene coexpression analysis (WGCNA), gene modules (rows) were identified which varied in their expression across the cell clusters (columns), shown here as a heatmap. The colours here signify the scaled average expression of gene modules across the various cell clusters. Gene modules were assessed for pathway enrichment (top six KEGG terms, Padj value < 0.05). The rightmost panel shows the number of genes within each gene module.

Of the 16 identified microglial cell clusters, four (cMG1-3, 6) expressed high levels of established homeostatic microglial genes, such as *Hexb, Tmem119*, *Gpr34* and *P2ry12* contained within gMG4, and in the absence of overt activation or cell cycle-related gene modules (see below). gMG4 was enriched for terms including P2Y receptors (*Padj* = 1.9×10^-2^) and TGF-beta receptor signalling (*Padj* = 7.4×10^-3^) consistent with a homeostatic state. cMG1 and cMG2 were predominant in CTRL samples and while there was overall similarity in profile between these clusters, expression of gMG7 genes enriched for lysosomal/phagocytic activity and ribosomal assembly/translation were greater in cMG2 (**Figure 4D**). cMG3 and cMG6 derived mostly from the MCAO condition and perhaps reflective of a minor deviation from the homeostatic state in response to injury, there was a mild reduction of gMG4 homeostatic genes and elevation of gMG7 lysosomal and ribosomal genes compared to cMG1/2 (**Figure 4D**).

The remaining microglial cell clusters expressed relatively lower levels of the homeostatic microglial gene set (gMG4) notably compared to cMG1-3 and cMG6 and were distinguished from each other by their induction of reactivity and/or cell cycle gene modules. Microglial cell clusters cMG7 and cMG16, both derived exclusively from the MCAO sample, expressed the lowest levels of the homeostatic gene set and shared marked induction of several large gene modules (gMG3, 5, 7) collectively comprising over 800 differentially-expressed genes enriched in multiple biological functions thus indicating the large-scale reactivity of these cells. High expression of gMG3 genes such as *Cst7*, *Spp1*, *Apoe*, *Lgals3* and various cathepsin genes (**Figure 4B-D**), indicated these cell clusters, particularly cMG7, as most resembling the DAM/ARM state first described in chronic cerebral proteinopathy models (we compare the stroke-associated microglial states with other disease contexts in-depth below). cMG7 and cMG16 also expressed high levels of other gene sets enriched in gMG3 involved in “lipid metabolism process” (e.g. *Apoe*, *Acadl*, and *Spp1; Padj = 8.50×10^-2^*), cell “oxidation-reduction process” (e.g. *Cyba/b*, *Cybb*, *Hmox1*, and *Sod2; Padj = 1.10×10^-13^*), and glycolytic and oxidative “ATP biosynthetic process” (e.g. *Pkm*, *and*, genes of mitochondrial complex; *Padj* = 1.74×10^-2^) (**Figure 4B-D**) implicating these microglial clusters as extremely metabolically active. In view of the influence of free iron on cell redox state, we noted with interest that expression of the genes (*Ftl1* and *Fth1*) encoding the iron storage protein, ferritin, was induced to high levels relatively specifically in cMG7 and cMG16 among all microglia (**Figure 4B**). The major difference distinguishing cMG7 and cMG16 was the high expression in cMG16 of cell cycle-enriched gene modules (gMG8, 11) (**Figure 4B**), perhaps indicating the positioning of cMG16 microglia at the transition between cell cycle and certain differentiated reactive states. cMG9, and to a lesser extent cMG4, both derived predominantly from MCAO samples, also showed induction of the reactive gMG3 gene module but of a weaker and less extensive nature compared to cMG7 and cMG16 (e.g. induction of the redox gene component was largely absent in cMG4 and cMG9), perhaps reflective of a transitional microglial state. However, cMG9 was notable for the highest *Apoe* expression of all microglia (**Figure 4B**). Among MCAO-enriched microglial clusters with suppressed homeostatic gene expression, cMG5 microglia were distinctive due to their high expression of interferon pathway genes, including *Irf7, Isg15, Stat1* and multiple genes from the *Ifi, Ifit* and *Ifitm* families (all contained within the gMG9 module enriched for ‘Interferon Signalling’ (*Padj* = 2.4×10^-3^)) (**Figure 4B-D**). cMG8 microglia were largely defined by their high expression of a small group of genes (modules gMG13 and more so gMG17), including *Ank*, *Axl*, *Cd9*, *Ccl6*, *Ccl9* and *Lpl*. While no microglial cluster showed a gene expression profile indicative of MHC-II antigen processing/presentation (discussed further below), cMG8 microglia were the only cells to mildly express genes such as *Itgax* and *Cd74*. cMG10 microglia shared similarities with cMG8 but were also distinguished by higher gMG13 module expression and their selectively high expression of chemokine and cytokine genes (gMG16: *Ccl3* and *Ccl4*, *Egr1, Il1b, Tnf*) (**Figure 4B-D**). In support gMG16 module signified by high expression in cMG10 was enriched in terms such as ‘Toll-like receptor signalling pathway’ (*Padj* = 1.6×10^-5^), ‘response to interleukin-1’ (*Padj* = 8.1×10^-4^) and ‘Chemokine receptors bind chemokines’ (*Padj* = 2.5×10^-2^) (**Figure 4D**). Interestingly, while cMG8 was overrepresented by cells from the MCAO condition (*P < 0.01*), cMG10 comprised of a significantly greater number of CTRL cells (*P < 0.05*), perhaps indicating an involvement of this state in microglial reactivity remote from the primary infarction as explored below.

### Microglial cell cycle trajectory induced by MCAO

Among microglia, several clusters (cMG11-16) expressed high levels of canonical cell cycle marker genes (e.g. *Pcna, Mki67, Top2a, Cdk1*) and WGCNA showed high expression of gene modules (e.g. gMG1, 2, and 12) enriched for cell cycle processes in these clusters (**Figure 4A-D, Table S3-S6**), consistent with our previous findings of proliferative microglia after experimental stroke^12^. We estimated the dynamics of this cell cycle response more thoroughly, first classifying all subclustered microglia according to cell cycle phase (G1, S, G2/M) using Seurat (**Figure 5A**) and visualising the expression pattern of the Seurat cell cycle gene set^45^, across cycling microglia and adjacent clusters (**Figure 5B**). Proliferative cells (S and G2/M) formed an arc (**Figure 5A**) in the UMAP and revealed a progression from S phase to G2/M gene expression. To further investigate these clusters and examine genes varying within this arc of proliferative cells, pseudotime analysis was conducted thereby modelling a trajectory from cMG11 > 13 > 14 > 15 > 12 > 16. The gene expression patterns were consistent with the modular gene expression pattern found using WGCNA. Notably, cMG13 and cMG14 microglia expressed high levels of G1/S transition and S phase genes, including *Cdca5*, *Ccne2,* and *Mcm6* whereas genes associated with G2/M, including *Cdc20, Ccnb2* and *Cenpa* were most highly expressed in cMG15 and/or cMG12 (**Figure 4B-D, 5C**). The frequency of G2/M annotated cells remained abundant although declined in cMG16 suggesting a continuum from cMG12 and towards exiting of the active cell cycle (**Figure 5B, C**). We noted with interest the rising expression in cMG16 of many genes that reached maximal expression in cMG7 reactive microglia across pseudotime (e.g. *Apoe, Flt1, Plin2, Spp1*) and that this coincided with the lowest expression across all clusters of homeostatic genes (e.g. *Cx3cr1, P2ry12, Hexb*) (**Figure 5C**). Coupled with the observation that cMG7 microglia (unlike other reactive state clusters) and cMG16 also clustered hierarchically by unbiased WGCNA (**Figure 4D**), this suggests potential close relationships between cell cycle and the emergence of selected forms of microglial reactivity.

**Figure 5.**
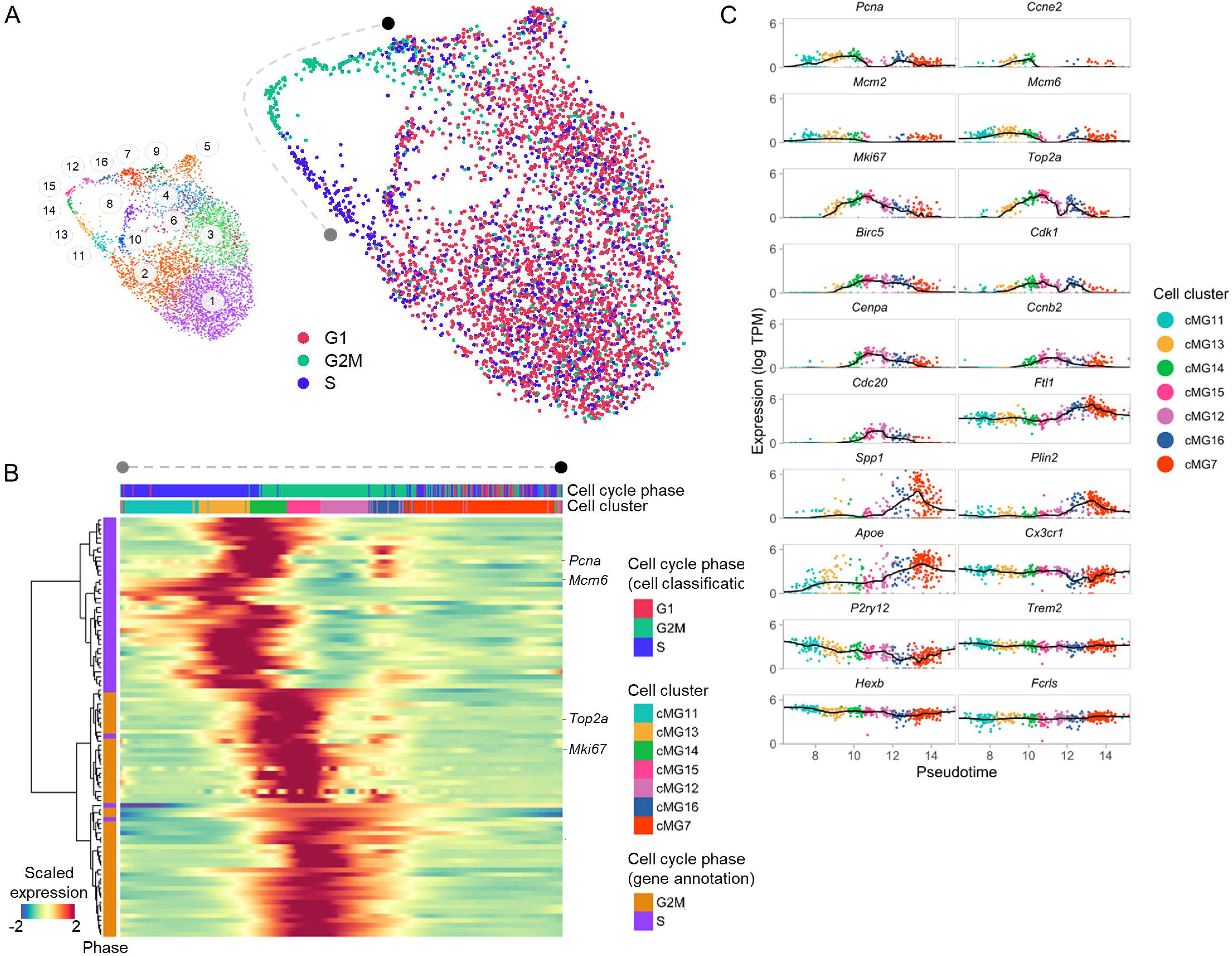
Microglial cell cycle analysis. (A) Cell cycle phase assignment of sub-clustered microglia using Seurat algorithm based on marker genes from each of the three phases. Colour-coded phase assignment is overlaid on the transcriptome-based clustered UMAP (inset) from **Figure 4**. Cells progressing through S to G2M forming an arc like arrangement are highlighted with endpoints marked in black and grey. (B) Heatmap of genes significantly (P< 0.05) changing along the arc of cycling cells. The genes (rows) are coloured based on their cell cycle phase annotation and the cells (columns) are coloured according to their assigned cell cycle phase as well as their transcriptome-based cell cluster. (C) Expression profile of selected genes representing cell cycle phases, microglial reactivity and homeostasis in cells from indicated clusters arranged along pseudotime. Each cell is coloured based on its transcriptome-based cluster membership (see **Figure 4** UMAP and inset in (A)).

### Comparative analysis of microglial states in stroke and other neurological conditions

The observations above indicated some microglial reactive states expressed gene modules resembling those described in other disease models. To formally examine this, we quantitatively assessed the correspondence between our stroke-associated microglial clusters and those in selected mouse models of neurodegenerative disease, ageing or inflammation for which scRNAseq datasets were accessible (**Figure 6, Table S7)**) ^46-53^. Briefly, the comparison was conducted by first acquiring differentially-expressed genes (DEGs) for the various microglial states across studies/models and measuring their degree of overlap with microglial cluster-defining DEGs identified from this study. The odds ratio was used as a measure of overlap and only significantly overlapping (odds ratio > 1 and *P* < 0.05) states were used in comparisons. Our stroke model cMG7 expression profile showed a substantial overlap with microglial clusters observed in most other conditions, including ageing, amyloidopathy, de/remyelination, and CNS trauma – this cluster comprises the state commonly termed “DAM/ARM” characterised by genes such as *Lgals3, Plin2*, and *Spp1*. The high odds ratio between our stroke model cMG5 and microglial clusters distinct from “DAM/ARM” observed in amyloidopathy (Sala-Frigerio, IRM)^51^, ageing (Hammond, OA3)^52^, neurodegeneration (Mathys, C6)^47^, and de/remyelination (Nugent, C8)^49^ was consistent with their signature of type 1 interferon pathway genes. These data indicate that IFN-enriched (e.g. IRM) and phagometabolic-enriched (e.g. DAM) microglial expression states are phenotypes common to multiple forms of CNS pathology, both acute and chronic. Contrasting with the above clusters showing widespread disease model intersection, the overlap between our stroke model cMG10 (enriched for chemokine and inflammatory genes such as *Ccl3, Ccl4 and Il1b*) and other models was more restricted, showing marked similarity only with ageing (Hammond, OA2)^52^ and CNS trauma (Milich, Inflammatory microglia)^48^. This may suggest this microglial expression state is associated with more specific types of injury/ageing-associated tissue damage or distress signals, although we do not exclude the possibility it could be detected at very early stages of a chronic disease process (not measured to date). Microglial proliferation is a feature of many CNS pathologies and the intersection of each of our stroke model cell cycle clusters (cMG12-16) with other acute (Milich, Inflammatory microglia)^48^ and chronic disease (Mathys, Cluster 3)^47^ model states is consistent with this. A particular observation of note was the marked overlap of our stroke model cMG16 cell cycle cluster with the TRM (which are likely transitioning to ARM) in CNS amyloidopathy^51^ but weak overlap with microglial clusters from other studies representing the IFN/ISG-enriched or chemokine-enriched states. As described above (**Figure 4, Figure S6**), cMG16 co-express certain gene modules with the ARM/DAM-like cMG7 after MCAO, their relatedness also highlighted by individual gene trajectories across pseudotime (**Figure 5**). In our cross-model comparison, cMG7 additionally overlaid markedly with the dividing microglial cluster in spinal cord trauma. Together, these data further implicate direct relationships between certain reactive microglial states and cell cycle that are relevant to multiple CNS pathologies.

**Figure 6.**
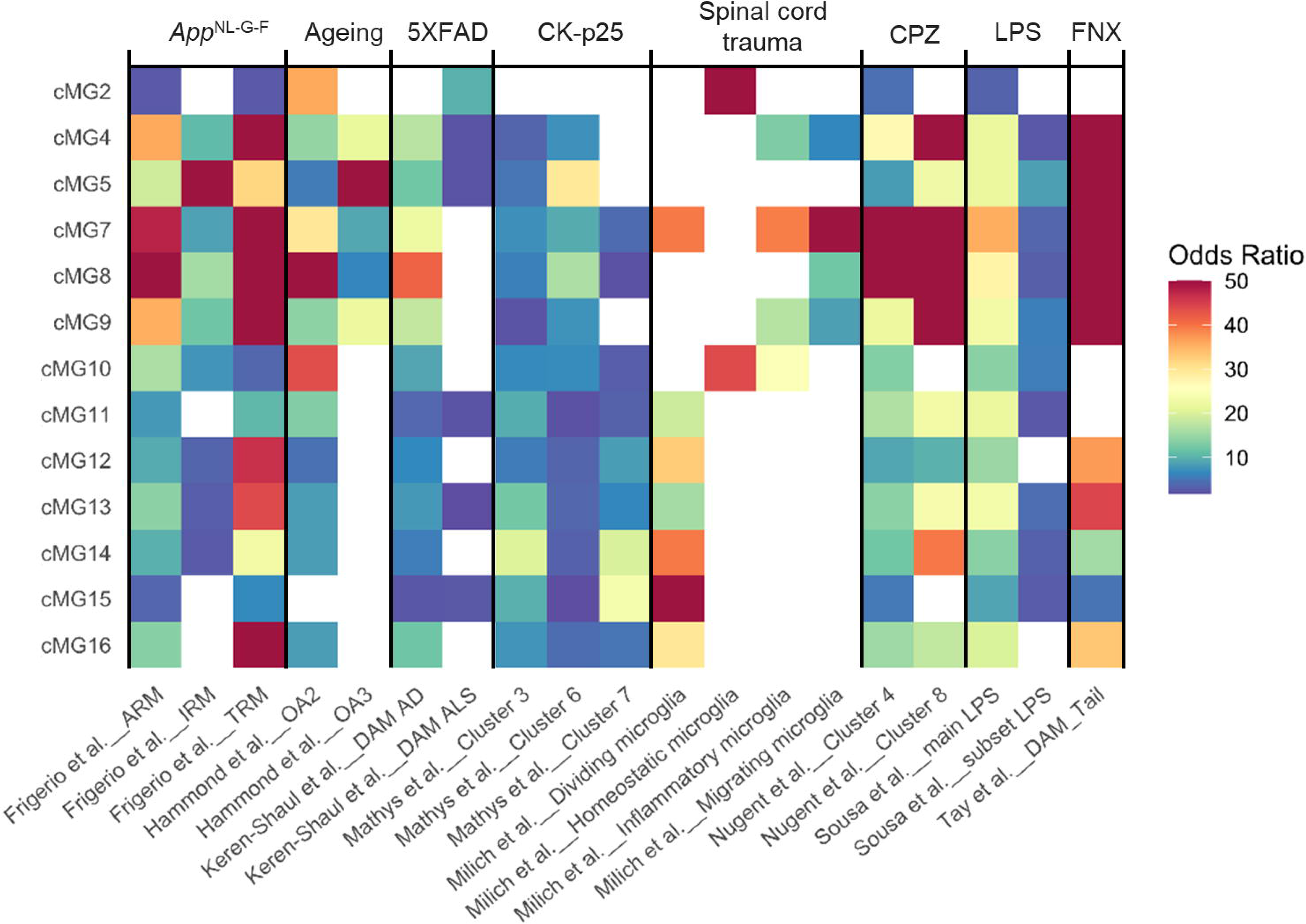
Cross-model comparison of microglial states. The top 500 differentially expressed genes across microglial clusters from this study (rows) were compared with selected clusters (see Methods) from other published datasets derived from models representing different disease, injury or neuroinflammatory conditions (columns) (see **Table S7** for studies used). Each comparison was made using the odds ratio which describes the degree of similarity (comparisons > 1 are more similar) and the significance of similarity given by a P value. Only significant comparisons have been shown where the odds ratio > 1 and P< 0.05.

### MCAO induces accumulation and differentiation of monocytes along two major trajectories

As described above, multiple myeloid cell clusters of non-microglial identity were evident, including CNS border-associated macrophages and several clusters of monocyte- and DC-related cells almost exclusively derived from MCAO samples (comprising almost one-fifth of all MCAO cells, **Figure 3D**). We specifically examined these cell populations by re-clustering all non-microglial (NMG) cells (excluding the small number of lymphocytes) and used differential gene expression (**Figure S7, Table S8**) and WGCNA (**Figure S8, Table S9**) to understand the cluster identities and characteristics (**Figure 7A-D, Table S10**). Two clusters were identified as CNS border-associated macrophages, their relatedness indicated by unsupervised hierarchical clustering of cell clusters according to gene co-expression across all non-microglial cells (**Figure 7D**). cNMG9 expressed high levels of the gNMG1 gene module genes including *Cd163*, *Clec10a* and *Vcam1* indicative of perivascular macrophages, whereas cNMG12 expressed relatively greater *Itgb5, Aif1* and MHC-II encoding genes, and a unique gene module (gNMG10) including *Axl*, with negligible *Clec10a* and *Cd163*, thus indicative of meningeal or choroid plexus macrophages^54^ (**Figure 7A-D**). cNMG5 cells expressed selectively high levels of *Nr4a1, Itgal,* and *Adgre4* (part of the gNMG8 gene module), and negligible *Ly6c2* and *Ccr2,* marking these as the patrolling/marginating Ly6C^lo/-^ monocytes. Conversely, cNMG4 cells expressed very high levels of *Ly6c2, Sell* and *Ccr2,* indicating these as stroke-induced Ly6C^hi^ monocyte immigrants given their almost exclusive derivation from MCAO samples (**Figure 7B**). cNMG7 retained *Ccr2* with diminished expression of gNMG7 module genes (e.g. *Ly6c2* and *Sell*), suggesting an early monocyte differentiation phenotype. Similarly, the gNMG7 gene set was diminished in cNMG3 cells but alongside induction of genes (mainly within the gNMG3 module) associated with cell-matrix adhesion and migration (e.g. *Ecm1*, *Fn1, Tgfbi*). gNMG3 gene module genes were highly induced in cNMG2 cells that additionally were defined by high expression of genes involved in macrophage lipid storage (*Plin2*), metabolism (*Lipa, Gpnmb*), lipoprotein assembly (*Apoe, Apoc1, Apobec1*) and cholesterol efflux (*Abca1, Npc2*). cNMG2 cells appeared in a highly transcriptionally active state given the extensive set of genes induced (e.g. gNMG3 module contains >500 genes, **Figure 7D, Figure S8**). Among several enriched functional gene classes, those involved in lysosomal acidification (*Atp6v* family) and enzymatic activity (including proteases, glycosidases, lipases, and nucleases) were most prominent. This enrichment was substantiated by KEGG analysis revealing “Lysosome” (*Padj* = 5.80×10^-8^) and “Phagosome” (*Padj* = 1.54×10^-4^) as enriched pathways (**Figure 7D**). We noted with interest that cNMG2 shared high expression of many genes (e.g. *Ctsb, Plin2, Spp1*) with certain reactive microglial clusters (**Figure 4**), a comparison we examine in more depth below. cNMG6 and cNMG10 cells shared high expression of a distinctive interferon response-enriched gNMG9 module (e.g. *Irf7, Ifit3, Isg15,* and *Oasl2*) with the distinction between these cell clusters the result of the additional expression of MHC-II antigen-presenting genes (e.g. *H2-Aa* and *H2-Eb1*) contained within the gNMG5 gene module in cNMG10 cells. This MHC-II peptide processing and presentation gene set was also highly expressed by cNMG1 cells regulator *Ciita* as one of the most highly induced genes in cNMG1, underlining their antigen-presenting cell phenotype and with high expression of *Flt3* and *CD209a*, but not *Zbtp46*, highly typical of a MoDC identity distinct from the cDC lineage^55^. *Zbtp46*, in contrast, was highly expressed with other canonical genes (e.g. *Ifi205*) as part of the gNMG6 gene module defining cNMG11 as conventional DCs^56,57^. cNMG8 cells were readily identifiable as migratory DCs by high expression of genes such as *Ccr7* and *Traf1*^56^.

**Figure 7.**
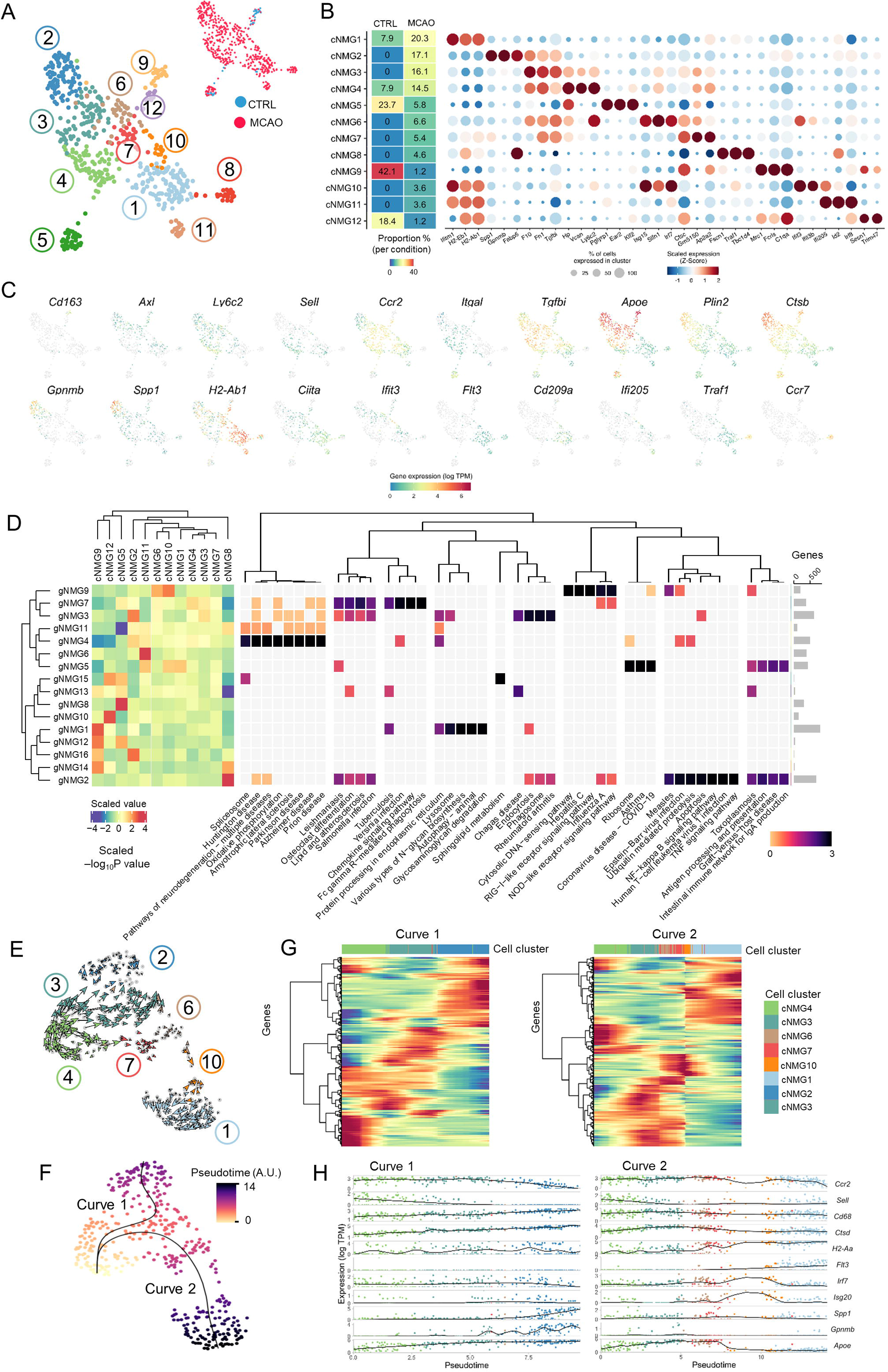
Heterogeneity and differentiation trajectories of non-microglial mononuclear myeloid cells associated with MCAO. (A) Non-microglial cells were sub-clustered from the entire dataset and based on their transcriptomic profile projected in 2D using UMAP. Inset UMAP shows the MCAO and CTRL conditions from which sub-clustered cells were derived from. (B) From left to right, for each cell cluster (rows) panels show the proportion of cells within the cell cluster derived from the MCAO or CTRL conditions, and a dot plot of the expression of marker genes. For the dot plot, the colours represent the mean expression value which is scaled across clusters and the size of the dot indicates the number of cells of the cluster (row) expressing each gene (column). (C) Expression of selected marker genes projected on the cell UMAP. (D) Using gene coexpression analysis (WGCNA), gene modules (rows) were identified which varied in their expression across the cell clusters (columns), shown here as a heatmap. The colors signify the scaled average expression of gene modules across the various cell clusters. Gene modules were assessed for pathway enrichment (top six KEGG terms, Padj value < 0.05). The rightmost panel shows the number of genes within each gene module. (E) RNA velocity analysis of cells from clusters annotated as Ly6C^hi^ monocytes and monocyte-derived cells (**Figure 7A-B; Table S10**) to estimate the directionality of altered gene expression inferring differentiation trajectory. RNA velocities are projected in 2D by UMAP. (F) Pseudotime analysis using Slingshot of the above Ly6C^hi^ monocytes and monocyte-derived cells. (G) Heatmaps for each curve from pseudotime analysis showing the genes (rows) which are altered significantly (P value < 0.05) across cells ordered along the trajectories. Genes are organised according to unsupervised hierarchical clustering which highlights the different patterns of gene coexpression associated with these trajectories. (H) Expression level (y-axis) of selected genes (dots, coloured by cell cluster) are shown along the pseudotime trajectories (x-axis).

The observations above implicated MCAO-induced differentiation of blood-derived *Ccr2*^hi^*Sell*^+^*Ly6c2*^+^ monocytes (cNMG4) towards highly differentiated monocyte-derived macrophage (cNMG2; enriched for lipid metabolism, lysosomal activity, matrix interactions) or monocyte-derived dendritic cell (cNMG1; enriched for MHC-II antigen presentation) fates. While cNMG1 and cNMG2 shared almost complete loss of *Sell* and diminished *Ly6c2* and *Ccr2* expression, entirely distinct transcriptional modules were induced (**Figure 7B-D, Table S8-S9)**. Potential transitional states were also evident, notably cNMG3 as an intermediary from monocytes to MDM and the interferon response-enriched cNMG10 transitioning towards MoDC. To explore these patterns more formally, we conducted trajectory inference, first using RNA Velocity (*Velocyto*)^58^ to determine the directionality of gene expression dynamics based on the ratio of unspliced to spliced mRNA (**Figure 7E**). This demonstrated a trajectory of transcriptional change progressing from cNMG4 (monocytes) towards cNMG2 (monocyte-derived or MDM) via cNMG3 and towards cNMG1 (MoDC) via cNMG7/6/10, thus consistent with the pattern of shared and distinct gene co-expression modules cross cell clusters above. We also conducted Pseudotime analysis^59^ to organise the cells along a pseudotemporal continuum (**Figure 7F**). Cells aligned along two curves representing a two-branched trajectory originating from cNMG4 (monocytes) that were abundant early in pseudotime in both curves and progressing along curve 1 towards cNMG2 (MDM) and along curve 2 towards cNMG1 (MoDC). Almost all the interferon response-enriched cNMG6/10 cells were positioned along curve 2, supporting their pre-MoDC expression profile, whereas cNMG3 cells were essentially the sole antecedent of MDM in curve 1 (**Figure 7F**). We determined the genes statistically highly-variable across pseudotime and grouped them according to their hierarchical clustering pattern (**Figure 7G**). This demonstrated the progressive gradients of expression both for gene sets induced and repressed along pseudotime with exemplar genes such as *Ccr2, Sell, Cd68*, and *H2-Aa* (**Figure 7H**) representative of the continuum in differentiation from monocyte to MDM and MoDC states. The transient induction of the interferon-response gene module mid-way along curve 2 was notable (**Figure 7G, H**) and further suggested these cell states as likely intermediaries preceding full acquisition of the MoDC phenotype.

### Comparative analysis of reactive microglia and monocyte-derived cells

The data above highlight distinct subsets/states within and between the microglial and monocyte-derived populations. Yet, despite the clear separation globally on UMAP between microglia and monocyte-derived clusters (**Figure 3**), similar gene expression traits were observed between these broad cell classes of distinct ontogeny in our analyses above. We were particularly interested in exploring more formally and at a cluster-specific level how the profiles of reactive microglia and the differentiated monocyte-derived cells related to each other, given their concurrent appearance and previous challenges in distinguishing these cells by bulk analyses. We integrated the sub-clustered microglial and non-microglial datasets (**Figure 8A**) and conducted WGCNA, thus generating an expression matrix of both shared and distinct gene modules across all mononuclear myeloid cell subclusters (**Figure 8B, S9, Table S12-13**). Myeloid gene modules (gMY) were annotated according to KEGG pathway analysis and manual inspection (**Figure 8B, Table S14**). Cell clusters formed three major top-level hierarchical groups comprising (i) all microglial states and BAM, (ii) Ly6C^hi^ monocytes and MDM, and (iii) cells of dendritic cell phenotype (including MoDC, cDC, MigDC) and Ly6C^lo^ monocytes. The DC grouping was most distinct among all cell clusters and was characterised as expected by high expression of antigen processing and presentation genes (gMY4: *Padj* = 3.27×10^-3^, gMY15: *Padj* = 1.22×10^-4^). Microglia did not express this antigen-presenting gene module whether of CTRL or MCAO sample origin, or of homeostatic or reactive transcriptional phenotype and more broadly, they shared few gene modules with the DC grouping, suggesting they retain highly distinct expression phenotypes from specialised antigen-presenting cells at this time point after acute ischaemic brain injury. In contrast, several microglial clusters co-expressed gene modules (gMY25 and gMY33) with the Ly6C^hi^ Mo/MDM grouping. gMY33 was enriched for lysosomal function (*Padj* = 5.27×10^-5^) and expressed by most microglial clusters, including those with moderate/high homeostatic gene expression (e.g. cMG1). However, the most striking similarity was evident between reactive MCAO-enriched cMG7 (DAM-like) microglia with severely repressed homeostatic gene set expression (gMY1: *Csf1r*, *P2ry12* and *Trem2*) and the highly differentiated MDM cluster (cNMG2). This relatedness was driven by co-expression of the above lysosomal gene modules (e.g. gMY29 containing *Cd68,* gMY16 containing *Ctsb*) with additional modules (gMY10, 23) that were notable for the presence of genes commonly considered part of the “DAM” microglial signature (e.g. *Spp1, Apoe*) and genes involved in glycolysis (e.g. gMG31 containing *Pkm; Padj* 2.17×10^-3^) (**Figure 8B-8C**). Alongside these shared characteristics, the transcriptional basis for the overarching distinction of these cell clusters was driven by gene modules reflecting the monocytic origin of MDMs (e.g. gMY17 containing *Ccr2, Sell*) and differences in oxidative metabolism and translational activity (e.g. gMY3: enriched for electron transport chain complex and mitoribosome genes). The gMY22 expression module comprised some genes expressed entirely selectively in MDM compared to cMG7 microglia, including *Arg1*, *Arg2 and Emp1* (**Figure 8D)**. We also noted several genes (e.g. *Lgals3, Plin2*) that, although expressed by both cMG7 microglia and cNMG2 MDM, levels were markedly greater and more consistently expressed across cells within the MDM cluster (**Figure 8E**). Differential gene expression analysis between these two specific clusters (cMG7 vs cNMG2) showed the individual genes that most differed (**Figure 8F, Table S15**).

**Figure 8.**
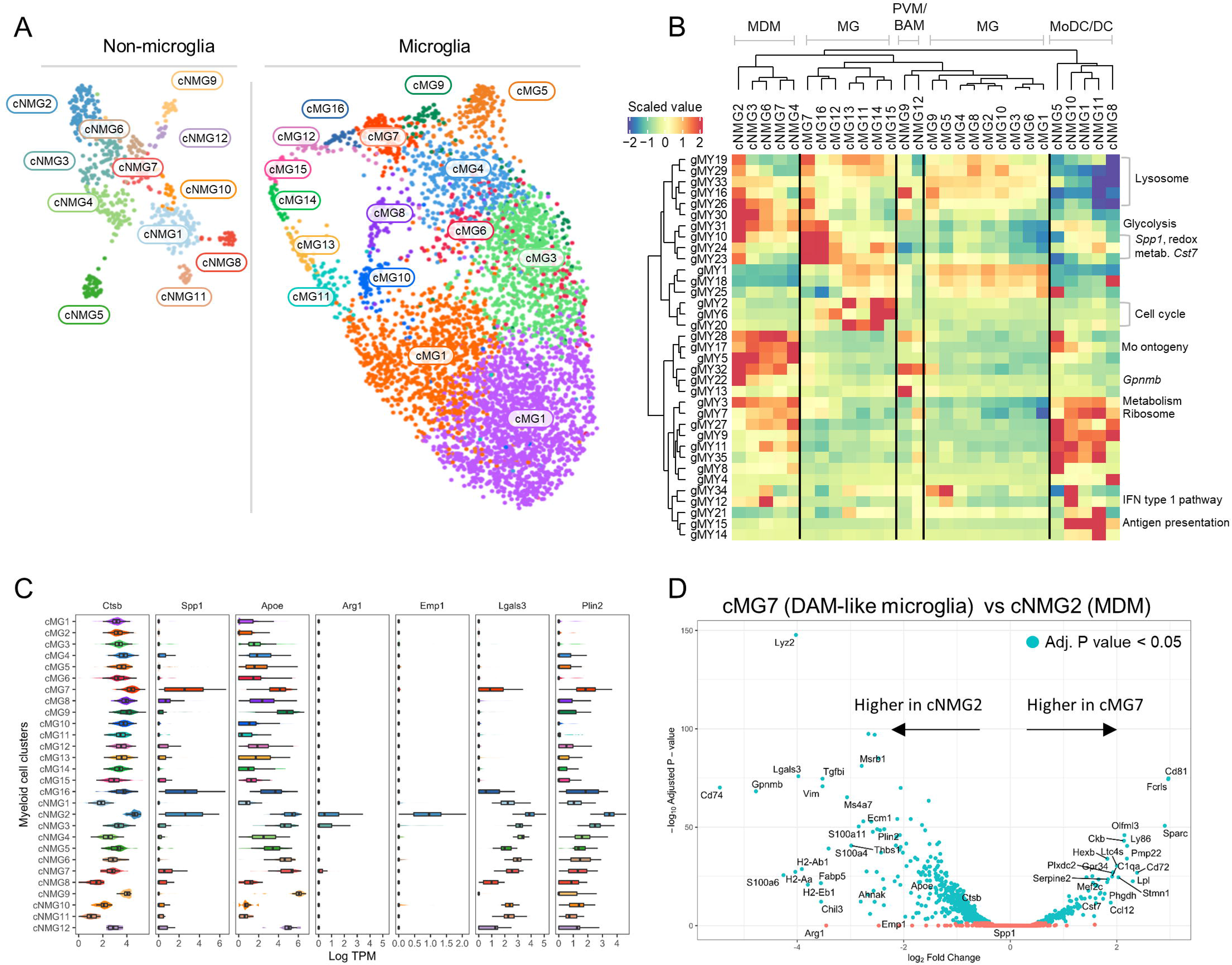
Comparative analysis of reactive microglia and non-microglial mononuclear myeloid cells associated with MCAO. (A) Composite transcriptome-based UMAP projection of sub-clustered microglia (right panel; from Figure 4A) and non-microglial myeloid cells (left panel; from Figure 7A). (B) Using gene coexpression analysis (WGCNA) gene modules (rows) were identified which varied in their expression across all the microglial and non-microglial myeloid cell clusters (columns), shown here as a heatmap. The colours signify the scaled average expression of gene modules across the various cell clusters. Gene modules (rows) and cell clusters (columns) are arranged according to unsupervised hierarchical clustering thus indicating inter-relatedness for gene modules and cell clusters. Major groupings of cell clusters and selected gene modules and representative genes are manually annotated for orientation (see Table S14 for full pathway analysis). (C) Expression distributions of representative genes (columns) across each microglial and non-microglial cell cluster (rows). (D) Analysis of differential gene expression between cMG7 (DAML) and cNMG2 (MDM). Differentially expressed genes (Padj < 0.05) are coloured blue and selected genes are highlighted. MDM, monocyte-derived macrophage; MG, microglia; PVM, perivascular macrophage; BAM, border-associated macrophage; Mo, monocyte; MoDC, monocyte-derived dendritic cell; DC, dendritic cell.

Overall, this comparative analysis conveys several key conclusions: (1) heterogeneity in myeloid cell transcriptional phenotypes 3 d after MCAO results from differences in combinatorial expression of multiple gene modules rather than all-or-nothing profiles, (2) comparisons of microglial and Mo/MdC transcriptional states at a bulk population-level obscure more fine-grained similarities and distinctions between cell subsets/states within these broad classes, (3) MCAO-enriched reactive microglial states generally show some transcriptional resemblance to MCAO-induced MDM, however none of the microglial clusters (reactive or steady-state) expressed antigen presentation gene modules exemplifying the distinct MCAO-induced MoDC/DC phenotypes, (4) greatest similarity is evident between an MCAO-enriched reactive microglial state with “DAM”-like properties and a highly differentiated MDM state, but these also retain distinction through differences in ontogeny- and metabolic-related expression modules.

### Integrated proteogenomic profiles of MCAO-associated myeloid cells

Our primary reason for using hashtag (antibody-conjugated) oligonucleotides (HTO) was to enable sample multiplexing and bio-replication (also assisting multiplet identification and removal (see above)). The HTO antibody detects the CD45 antigen hence a further application of this method is to use sequencing output from the HTO as a measure of cell surface CD45 expression, which can then be integrated with transcriptome profiles on individual cells, a proteogenomic approach called CITE-seq^43^. As described above, surface CD45 levels on flow cytometry are often used to broadly distinguish microglial and non-resident brain mononuclear myeloid populations for analysis or isolation, however the relationship between CD45 intensity and subpopulation phenotypes, including their transcriptional state, has not been possible to study previously. We reasoned that a proteogenomic approach using HTOseq output for surface CD45 overlaid on the mRNAseq-defined myeloid cell clusters (**Figure 8**) could explore this. The range of HTOseq intensity distributions was broad across the entire set of myeloid cell clusters, ranging from lowest in cMG1 and cMG2 (homeostatic microglia) to highest in cNMG7 (monocytes) and cNMG2 (MDM) (**Figure S10A**). Ordering of cell clusters according to HTOseq median intensity showed all monocyte-derived clusters above all microglia. Data indicated that for HTOseq intensities representing even the most highly expressing microglial clusters, the probability of mis-classifying a cell as monocyte-derived was low thus supporting the general distinction (CD45^hi^ vs CD45^lo^ for the parent populations) commonly used in conventional cytometry gating. PVM/BAM showed an intermediate HTOseq intensity (**Figure S10A**) in keeping with known higher CD45 protein measured by flow cytometry compared to steady-state parenchymal microglia^60^. However, how levels compare to reactive microglia in stroke brain is unknown. Our data show close overlap with highly reactive microglia (e.g. cMG7) underlining the importance of using PVM/BAM protein markers in stroke cytometry studies. Within microglia, the greatest HTOseq intensities were evident on the cell cycle and DAM-like clusters (with cMG7 and cMG9 highest of all microglia), providing orthogonal evidence for these microglial states as the most highly reactive (**Figure S10A-B**). Transgene-based reporters, fate mapping, scRNAseq and combinations thereof will remain the gold standards for cell ontogeny-phenotype determination, however these are resource-heavy approaches. The present data, while derived from a single cell surface protein HTO barcode, show the principle for how a proteogenomic approach could be used to identify widely deployable low-plex surface protein antibody combinations with potential to report on -omics-defined intracellular states thus extending the capabilities of existing conventional cytometry in future stroke studies.

### *In situ* analysis reveals spatial organisation of myeloid cell responses associated with structural brain connectivity after MCAO

By sampling cells from hemispheres ipsilateral and contralateral to arterial occlusion, our scRNAseq expression patterns implicated spatially-organised cell types and states. We used multiplexed single molecule detection FISH to corroborate and extend these findings, in particular to examine specific brain areas near and remote to the infarct. During our single-cell analysis of microglial clusters in MCAO tissue (**Figure 4**), we noted the appearance of some cell clusters (cMG3, 4, 5, 7, 8, 12 and 16) predominantly in ipsilateral hemispheres, whereas others were also present in contralateral hemispheres. To orthogonally validate this spatial heterogeneity, we chose to compare the regional distribution of cMG7 (DAM-like), which was exclusively found in ipsilateral hemispheres, with cMG10 (chemokine-enriched), which was found in both, using the cluster-defining genes *Spp1* and *Ccl3* alongside the microglial/BAM-specific marker *Fcrls* (**Figure 9A - B**). It was first evident that *Ccl3* was not expressed at detectable levels in brain tissue in sham-operated controls, whereas, both *Fcrls* and *Spp1* were (**Figure 9A**). *Fcrls* was uniformly spaced throughout the brain (consistent with the tiled distribution of microglia), whereas *Spp1*^+^ puncta were observed in the cortex, striatum and septal nuclei. High-magnification confocal imaging of combined *Fcrls, Ccl3* and *Spp1* revealed *Fcrls* did not colocalise with either cluster-defining marker in sham control brains, indicating both phenotypes were reactive to MCAO and not features of a homeostatic or surgically-induced microglial landscape (**Figure S11** and **Figure 9B**). In contrast, labelling of all three genes was increased in the brain 3 d post-MCAO, albeit with differing spatial distributions (**Figure 9A**). The increased labelling of both *Fcrls* and *Ccl3* was observed in peri-infarct regions and along brain regions structurally connected to the cortical infarct/peri-infarct, such as transcallosal and cortico-striatal fibres. On the contrary, MCAO-specific *Spp1* labelling was highly restricted to infarcted and peri-infarct brain tissue (**Figure 9A-B**), consistent with our previous finding that cMG7 was specifically isolated from ipsilateral hemispheres (**Figure 4B**). Quantification of high-magnification confocal imaging confirmed presence of *Fcrls*^+^*Ccl3*^+^ and *Fcrls*^+^*Spp1*^+^ cells around the infarct that were absent in sham controls (*Ccl3*: *Padj =* 8.5×10^-14^, *Spp1: Padj =* 1.1×10^-12^). *Fcrls*^+^*Ccl3*^+^ cells were, however, ∼10-fold more abundant than *Fcrls*^+^*Spp1*^+^ cells along transcallosal fibres in the corpus callosum (*Padj =* 2.4×10^-6^, **Figure 9B**) and ∼6-fold in the contralateral cortex (*Padj* = 0.002, **Figure 9B**). Thus, *Ccl3*^+^ microglia (i.e. reflective of cMG10) appear in/around both primary infarcted tissue and in perturbed communicating fibres in remote structurally connected brain regions. *Spp1*^+^ microglia (i.e. reflective of cMG7), on the other hand, are mostly restricted to infarct/peri-infarct tissue 3 d post-MCAO. However, a small but significant increase in cell number above control tissue in the corpus callosum (**Figure 9B**, *Padj* = 0.001) may indicate this reactive state also appears in connected regions.

**Figure 9.**
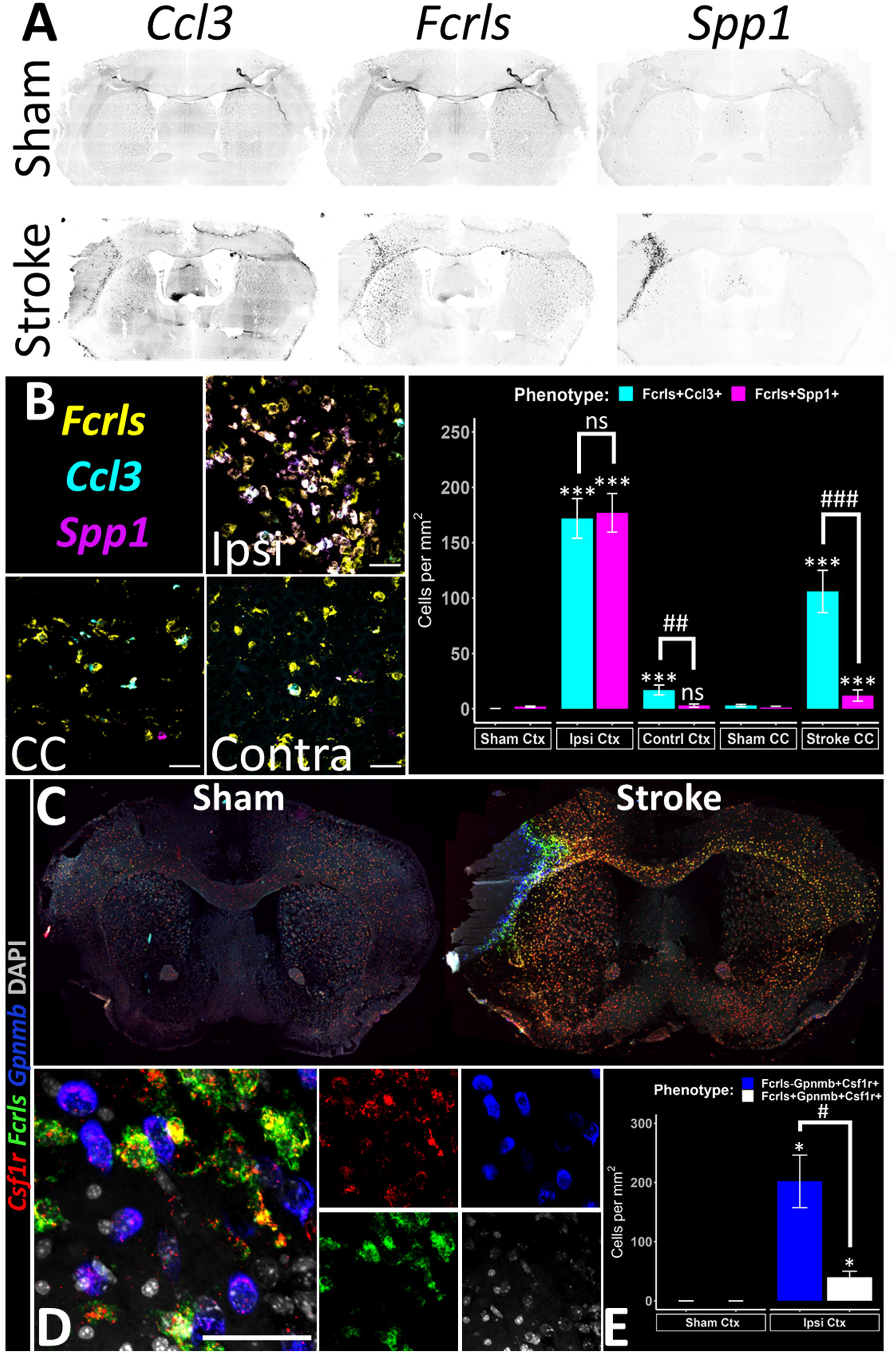
Spatially resolving microglial and non-microglial transcriptional phenotypes. (A) Slide-scanned images of multiplex fluorescent in-situ hybridisation for transcripts Ccl3, Fcrls and Spp1 in (top) Sham-operated control murine brains and (bottom) three days following unilateral middle cerebral artery occlusion (Stroke). Individual channels have been separated and presented in grayscale. (B) Maximum intensity projections from high-power confocal stacks in ipsilateral peri-infarct (Ipsi), Corpus Callosum (CC) and contralateral Cortex (Contra) of Stroke brains (scale bar = 50µm). (iv) Fcrls^+^ Ccl3^+^ (cyan) and Fcrls^+^ Spp1^+^ (magenta) phenotypes were manually quantified from 3 – 6 non-overlapping regions of Sham (F**igure S11**) and Stroke (B) and abundance compared across brain regions (linear mixed modelling with Holm-Sidak posthoc pairwise comparisons; * = significantly different to sham and # = significantly different to other phenotype). (C) Slide-scanned images of multiplex fluorescent in-situ hybridisation for transcripts Csf1r, Fcrls and Gpnmb in Sham and Stroke. (D) Maximum intensity projection of transcript expression in the peri-infarct region with (left) combined and (right) individual channels shown (scale bar = 50µm). (E) Fcrls-Gpnmb^+^ Csf1r^+^ (blue) and Fcrls^+^ Gpnmb^+^ Csf1r^+^ (white) phenotypes were manually quantified in Sham cortex (left) and peri-infarct of Stroke (right) and cell abundance compared across phenotypes (* = significantly different in one sided t-test against 0; # = significantly different in paired t-test; bonferroni correction applied for 3 comparisons). ns = not significant, * = padj<0.05, ** = padj <0.01, *** = padj <0.001; n=4 per group. Data expressed as mean ± SEM.

*Spp1* was highly enriched in both cMG7 (DAM-like) and cNMG2 (MDM) in our previous single cell analyses (**Figure 4** and **Figure 7**) and thus likely labelling cells of both ontogenies *in situ*. To better separate these two cell classes, we performed smFISH on the pan-mononuclear phagocyte marker *Csf1r*, cNMG2-specific marker *Gpnmb* and microglial/BAM-enriched *Fcrls* (**Figure 9C – E**). *Gpnmb* was notably absent in sham control brain tissue, whereas *Csf1r* and *Fcrls* colocalised across the entire brain section (**Figure 9C**). In contrast, *Gpnmb* was strongly expressed within and around the infarct of MCAO brain tissue as was *Fcrls*, although, the latter was upregulated along connected brain regions as previously noted. High-magnification confocal imaging revealed *Gpnmb*^+^ cells expressed low levels of *Csf1r* compared to neighbouring *Fcrls*^+^ cells (**Figure 9D**), consistent with the finding that *Csf1r* is more highly expressed by microglia and BAMs over cells of bone marrow origin (**Figure 3E**). These *Gpnmb*^+^*Fcrls*^-^*Csf1r*^+^ (cNMG2, MDM) cells were ∼5-fold more abundant in peri-infarct tissue than *Gpnmb*^+^*Fcrls*^+^*Csf1r*^+^ cells, which likely represent the very small amount of reactive microglia expressing *Gpnmb* (**Figure 9E**, *Padj* = 0.04). Together, these data indicate cNMG2 MDM recruited to the brain following MCAO are confined to the infarct and peri-infarct tissue, in close proximity to cMG7 microglia that share many transcriptional features.

Our earlier single cell analysis indicated there were two potential routes to increasing the brain macrophage pool following stroke, dependent on cell ontogeny. One route utilised the unique ability of microglia, as the tissue-resident macrophage population, to proliferate whilst the other relied on recruitment of immature cNMG4 monocytes and subsequent maturation into cNMG2 MDMs (**Figure 3, 5, 7**). To spatially resolve these differences, we once again performed smFISH using *Fcrls*, the monocyte marker *Ccr2* and the cell cycle marker *Mki67* (**Figure 10A-D**). Labelling of *Ccr2* was notably absent in sham-operated controls and *Mki67* labelling in shams was consistent with known anatomical proliferative cell niches in steady-state (periventricular zones and hippocampus) (**Figure 10A**). *Mki67* rarely colocalised with the microglia/BAM marker *Fcrls* in control brains (**Figure 10B**). 3 d post-MCAO, *Ccr2* was present within and around the infarct and *Mki67* was found in peri-infarct regions and in structurally connected regions, such as the thalamus (**Figure 10A**). High-magnification confocal microscopy revealed ∼15% of the *Fcrls*^+^ cells (∼50 of a total ∼350 *Fcrls*^+^ cell/mm^2^ pool) in peri-infarct tissue were positive for the cell cycle marker *Mki67*, whereas none of the *Ccr2*^+^ cells were *Mki67*^+^, despite the close proximity of *Fcrls*^+^ and *Ccr2*^+^ cells (**Figure 10B-C**). No *Ccr2*^+^ cells were observed in the corpus callosum or contralateral cortex of MCAO brains, although, some *Mki67*^+^*Fcrls*^+^ cells could be seen in these regions structurally connected to the primary infarct (**Figure 10D**), consistent with the respective unilateral and bilateral recovery of these cells in our single cell experiment (**Figure 3**). We next looked to compare *in situ* labelling of the *Mki67* gene with immunolabelling for the encoded protein Ki-67 (**Figure 10E-G**). In line with transcript-level data, Ki-67^+^P2Y12^+^ microglia were rare in sham-operated control cortical brain tissue (**Figure 10E**). However, a large amount of Ki-67^+^P2Y12^+^ microglia were evident in peri-infarct tissue 3 d post-MCAO. Most of these Ki-67^+^ microglia were more amoeboid and expressed lower levels of P2Y12 than neighbouring Ki-67^-^ ramified microglia that were a greater distance from the infarct border. Such reduction in P2Y12 expression is consistent with our transcriptional dataset indicating microglia downregulation of homeostatic markers is maximal in cycling microglia (**Figure 5C**). We were also able to visualise the appearance of stroke-specific Ki-67^+^P2Y12^+^ hypertrophic microglia in the corpus callosum (**Figure 10F**) and contralateral cortex (**Figure 10G**). Together, these data indicate microglia enter the cell cycle around infarcted tissue and in structurally connected remote brain regions, and downregulation of homeostatic markers and retraction of cell processes are associated with this. We also note with interest the close proximity of proliferating microglia, immigrant monocytes and differentiated MDMs in peri-infarct zones, which implies shared molecular determinants of mononuclear phagocyte fate at this timepoint after stroke independent of ontogeny.

**Figure 10.**
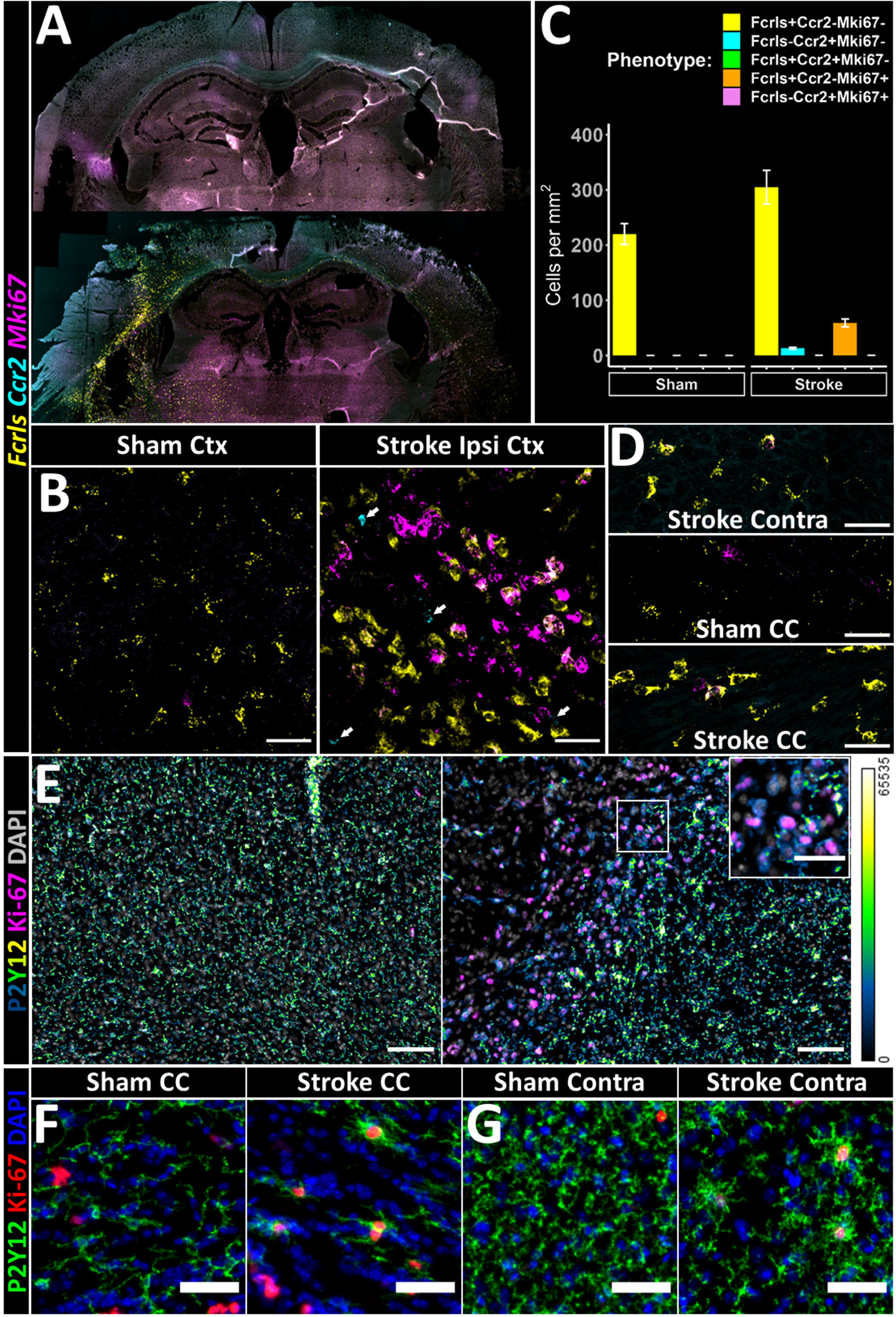
Spatially mapping cell cycle microglia at transcript and protein level. (A) Slide-scanned images of multiplex fluorescent in-situ hybridisation for transcripts Fcrls, Ccr2 and Mki67 in (top) Sham-operated control murine brains and (bottom) three days following unilateral middle cerebral artery occlusion. (B) Maximum intensity projection from high-power confocal stacks in (left) Sham Cortex and (right) Stroke peri-infarct. Ccr2^+^ monocytes (arrows) can be observed in peri-infarct regions in close proximity to Fcrls^+^ Mki67^+^ reactive microglia. Scale bar = 50µm. (C) Fcrls^+^ Ccr2- Mki67- (yellow), Fcrls- Ccr2^+^ Mki67- (cyan), Fcrls^+^ Ccr2^+^ Mki67- (green), Fcrls^+^ Ccr2- Mki67^+^ (orange) and Fcrls- Ccr2^+^ Mki67^+^ (violet) phenotypes were manually quantified from 3 – 6 non-overlapping regions of Sham Cortex and Stroke peri-infarct (B). (D) Representative images of Stroke contralateral Cortex (Contra), Sham Corpus Callosum (CC) and Stroke CC scale bar = 50µm. (E) Protein Ki-67 immunolabelled in (left) Sham Cortex and (right) Stroke peri-infarct alongside the homeostatic microglial marker P2Y12. P2Y12 intensity is visualised on a continuous scale with black/blue representing low, green average and yellow/white high expression (scale right). Inset is higher resolution of white box. Scale bar large image = 100µm, inset = 50µm. (F) Representative images of Ki-67 and P2Y12 labelling in the Corpus Callosum of (left) Sham and (right) Stroke and (G) contralateral Cortex of same groups, scale bar = 25µm. n=4 per group. Data expressed as mean ± SEM. 2

## Discussion

The present study brings new insight to the spatial organisation and composition of the mononuclear myeloid cell response to experimental stroke at a crucial transitional timepoint from acute to sub-acute phases. Markedly expanded cellular diversity within the overall mononuclear myeloid cell population was evident in MCAO samples. Within microglia, the relative proportion of the main homeostatic state cluster (cMG1) declined to less than 15% of cells in the MCAO condition. By cross-referencing our scRNAseq cluster proportions and quantitative cytometry data, we can estimate that despite the number of total microglia doubling in MCAO samples the absolute number of homeostatic state microglia declined by half, suggesting that state conversion of microglia already *in situ* as well as *de novo* microglial production via proliferation generates this more abundant and more diverse microglial population. The monocyte-derived cell clusters were almost exclusively derived from MCAO samples indicating that, as expected, they are adding tens of thousands of newly recruited cells to supplement the local CNS myeloid pool. The need for this increased cellular diversity presumably reflects the array of *de novo* environmental signals and complex tissue demands requiring a greater range of cell specialisations in the aftermath of stroke, particularly at this time point as tissue repair processes are triggered by an initial wave of damage and danger/distress signals^4^.

It is well established that microglia alter their transcriptome at a population level following acute stroke^14,28^ and recent preclinical studies have shown important functional consequences of global microglial elimination on stroke injury and repair^16,20,61,62^. However, reactive heterogeneity within the overall microglial population and their spatial organisation are poorly understood. Excluding microglia in cell cycle, our scRNAseq data showed multiple microglial clusters with marked suppression of a panel of canonical homeostatic genes and/or induction of separate gene modules expressed at negligible levels in homeostatic microglia. These differed qualitatively according to their combinations of elevated metabolic, inflammatory and phagolysosomal gene modules, but also quantitatively by the size of the gene networks involved, evident through the gene coexpression analysis on sub-clustered microglia (**Figure S6**). The cMG7 microglial cluster was notable for its vast array of genes altered, many associated with bioenergetic, redox and lipid metabolic processes marking this state as particularly reactive. cMG5 (type 1 IFN-enriched) and cMG10 (chemokine-enriched), in contrast, highly elevated entirely distinct gene sets of smaller size. This might suggest segregation of functional properties perhaps through exposure to specific stimuli. This is supported by striking spatially-dependent expression of microglial transcripts characteristic of these distinct scRNAseq clusters revealed by multiplexed ISH. In close agreement with scRNAseq showing cMG7 derived largely from samples ipsilateral to MCAO, *Spp1*^+^*Fcrls*^+^ microglia were most abundant in the peri-infarct area and elevated in number compared to sham-operated controls. In contrast, scRNAseq cMG10 cells derived more equally from contralateral and ipsilateral samples, which was matched by the detection of *Ccl3*^+^*Fcrls*^+^ microglia both in peri-infarct areas and remotely, including in contralateral regions. Beyond this bi-hemispheric pattern, there was a non-random distribution of these contralateral *Ccl3*^+^*Fcrls*^+^ microglia with most located in cortical areas connected by axonal tracts to the infarct and peri-infarct region, and along the axonal tracts themselves (e.g. callosal fibres). This specific pattern, and the elevated numbers of *Ccl3*^+^*Fcrls*^+^ microglia in MCAO compared to sham-operated mice also supports that these remote reactive microglia and the cMG10 cluster are caused by induction of MCAO and not surgical stress or reflecting a pre-existent baseline microglial state. Given their anatomical distribution, this state of microglial reactivity may be associated with altered structural or functional neural connectivity between regions proximal and distant to the primary stroke injury. Remote neurodegeneration and neuroplasticity in connected brain areas are increasingly recognised to occur after acute stroke and are implicated in long-term motor and cognitive outcomes^33,35,39^. The presence of morphologically-reactive microglia in areas of secondary neurodegeneration has been reported previously^37^. Our data now show the molecular signatures characteristic of remote reactive microglia that arise early after stroke and in the absence of overt secondary degeneration, perhaps suggesting more acutely altered neuroaxonal signalling as a trigger.

Recent studies using scRNAseq, mostly conducted in chronic disease models and ageing, have identified microglial transcriptional profiles that emerge in these various conditions^63^, including some that appear relatively well conserved across different chronic disease pathologies. Understanding the pathological conditions that share common reactive microglial states and those associated with more distinct profiles is likely to offer clues to the specific molecular cues that guide the emergence of different reactive states. Our quantitative analysis showed that MCAO-enriched cMG7 microglia shared significant similarity with those observed in CNS amyloidopathy, chronic de/remyelination, spinal cord trauma, axotomy, and ageing (**Figure 6**). Their common profile is defined by high expression of marker genes such as *Lgals3, Spp1, Plin2*, and *Apoe*, multiple elevated cathepsin and metabolic genes, an expression state originally referred to as the DAM/ARM state upon their discovery in CNS amyloidopathy models^46,51^. Two recent studies using a different (transient) stroke model also observed a cluster of microglia/myeloid cells with the characteristic DAM/ARM profile^10,30^ and while some stroke-specific elements have been proposed there appear many more similarities than differences with chronic disease DAM/ARM. The induction of a microglial state recently demonstrated after spinal cord trauma (termed “Migrating microglia”, see Fig 6)^48^ is largely indistinguishable from the profile we show here after MCAO and this spinal trauma phenotype also overlapped closely with chronic disease microglia. In both stroke and spinal cord trauma the DAM-like state is evident within days of the insult, highlighting that this expression state of microglia is not confined to chronic disease but is a conserved response evident in multiple acute and chronic pathologies. Whether cells converge on to this expression state despite exposure to different signals or there is an as-yet unidentified signal common to all pathologies remains to be determined. As noted above, *Spp1*^+^*Fcrls*^+^ microglia were mostly located in peri-infarct areas after MCAO suggesting that signals predominating in this tissue environment favour a DAM-like phenotype. The *Spp1*^+^ cMG7 microglial cluster we describe here expressed high levels of lipometabolism genes such as *Apoe, Fabp5, Nceh1*, and *Plin2*. Debris and lipid processing are a feature of peri-infarct remodelling and our data align with the recent report of lipid-laden and lipid-processing microglia/macrophages of similar transcriptional profile in the infarcted tissue after transient MCAO^30^. Our cross-model comparison also clearly showed that our MCAO-associated cMG5 profile was shared by clusters in other conditions (proteinopathies, ageing, nerve injury) but was distinct from the cMG7/DAM overlap pattern and defined by its type 1 interferon pathway-enriched profile. Therefore, this IFN/ISG-enriched reactive state also appears relatively conserved across pathological conditions. Recent studies have begun to indicate important disease-modifying roles for IFN-enriched microglia in chronic disease and ageing^52,64^. In contrast to the above commonly shared states, the CCL chemokine-expressing cMG10 profile was relatively less well represented across disease conditions, although evident in ageing and spinal cord injury. This state may be associated with signals more prevalent in acute than chronic disease, further supported by its presence in acute^52^ but not chronic^49^ demyelination models, although we do not exclude that a microglial state with this profile may arise in earlier phases (not yet studied) of chronic disease. Given the *Ccl3*^+^*Fcrls*^+^ microglia after MCAO were associated spatially with white matter tracts and brain areas remote but connected to the primary infarct, it is tempting to speculate that this state may be driven by altered neuroaxonal activity and/or glia-axonal signalling prevalent in areas distant from the primary injury. Further studies are needed to uncover the regulatory and functional attributes of this and other reactive microglial states, including if these discrete expression states map to correspondingly distinct functional states with regional effects.

We detected several microglial clusters with a cell cycle signature, amounting to ∼9% of MCAO-derived microglia, and *in situ* analyses showed ∼15% of peri-infarct microglia were Ki67^+^. This is 10-15-fold greater than basal microglial proliferation rate underpinning steady-state population self-renewal^65^. scRNAseq and smFISH demonstrated that cell cycle-engaged *Mki67^+^Fcrls^+^* microglia were mostly MCAO hemisphere-derived and localised to the peri-infarct zone. This might suggest that release of tissue damage molecular signals and/or the altered cellular milieu within the peri-infarct zone is a particularly strong trigger for MCAO-associated microglial proliferation. Our data clearly show that microglia but not Mo/MdC are in cell cycle at this timepoint, consistent with a recent scRNAseq study in spinal cord trauma^48^. This is despite the close spatial intermixing of microglia and MDM in peri-infarct tissue and ostensibly similar environmental exposure, thus suggesting injury-induced cues combine with ontogeny-determined factors to influence cell cycle entry. Inhibiting microglial proliferation after MCAO is detrimental to outcome^66^ whereas protective effects have been reported in chronic disease settings^67-69^. This highlights that while microglial proliferation is a common feature of most CNS disease and potential target for manipulation, the functional role of proliferation beyond population expansion is relatively poorly understood. Our data here point to a potentially important relationship between cell cycle and microglial reactive state development. Coexpression analysis and pseudotemporal ordering both suggested the G2M phase-enriched cMG16 as an intermediary state between cell cycle and the DAM-like cMG7. WGCNA of differentially-expressed genes showed several gene modules shared by cMG16 and cMG7 resulting in their unsupervised hierarchical clustering thus emphasising their transcriptional relatedness. In contrast, other reactive states did not show this connection with cycling microglia. Moreover, pseudotime analysis showed a progressive decline in microglial homeostatic genes (e.g. *Cx3cr1, P2ry12*) through cell cycle phases in parallel to rising expression of characteristic DAM/ARM genes such as *Apoe, Plin2*, and *Spp1*, notably at the cMG16/cMG7 transition. CSF1R and TREM2 are both known regulators of microglial proliferation. CSF1R inhibitors lowered whereas TREM2 agonists augmented the frequency of DAM/ARM-like cells within the microglial population during CNS amyloidosis^70,71^. Microglia with highly overlapping phenotype to this DAM/ARM-like state, that we show here to be conserved across multiple neuropathologic conditions, are also transiently present in the early postnatal brain in highly proliferative niches associated with normal developmental ^72^. Collectively, these data warrant that future studies more deeply explore the connections between microglial cell cycle and emergence of reactive states.

The influx of monocyte-derived cells to the post-stroke brain is well established and our scRNAseq data highlight that they contribute >15% of the cells forming the markedly more abundant and phenotypically diverse mononuclear myeloid population in MCAO. A key function of injury-recruited monocytes is to differentiate to more specialised derivatives which after stroke are not well defined. Our clustering and pseudotemporal analyses show clear differentiation along two major pathways, one producing cells with a highly specialised macrophage phenotype enriched in phagolysosomal and lipometabolism gene modules, and the other generating cells of dendritic cell phenotype highly enriched for antigen presentation genes. Importantly, we observed that these differentiated cells and the preceding intermediary phenotypes all emerged from Ly6C^hi^ monocyte precursors indicating that in the acute/subacute period Ly6C^hi^ but not Ly6C^lo^ monocytes are the sole source for these non-resident differentiating MDM and MoDC. This dual trajectory of differentiation solely from Ly6C^hi^ monocytes has also been reported recently in cardiac ischaemia^73^. At later time points after stroke it is unclear if Ly6C^lo^ monocytes may contribute although this appears unlikely in most tissue injury contexts^13,74^. It is intriguing to consider if the differentiation fates of monocytes are dictated stochastically upon arrival in the injured brain or there is already some pre-restriction of each cell to certain paths as has been proposed during haematopoietic development^75^. We noted with interest that the MoDC cluster, exclusive to the MCAO condition, was the most abundant that demonstrated a DC phenotype. This MoDC cluster, alongside cDC clusters, but not microglia, highly expressed a module of antigen processing/presenting and costimulatory genes (e.g. *Cd74, H2-Aa, H2-Ab1, Dpp, Btla*) including the key regulator *Ciita*. The complete segregation of MoDC and cDC clusters from all microglia by unsupervised clustering of cell clusters based on gene module coexpression reinforces the lack of any DC-like properties of microglia at this timepoint. Despite low level *Itgax* expression in a small number of MCAO-associated microglia, these data implicate infiltrating myeloid cells, rather than microglia, as the major antigen presenting cells. In support, a recent study showed that brain infiltrating myeloid cells are functionally much more capable than microglia of stimulating T cell proliferation after MCAO and that CD11c expression on microglia does not correlate with any difference in their antigen presenting capacity^76^. In contrast to the distinctiveness of all microglial clusters and MoDC, our unbiased WGCNA-based clustering of all myeloid cell clusters showed a striking similarity between the MDM (cNMG2) and DAM/ARM-like microglial (cMG7) clusters based on shared high expression of multiple gene modules. These contained many archetypal DAM/ARM genes and were also enriched for lysosomal, phagocytic and lipometabolic processes. The functional pathways represented by this multi-module overlap can be considered to involve “effector” elements of microglial/macrophage identity and function, and explain how it has been challenging to distinguish between resident microglia and non-resident macrophages when using individual markers associated with these effector functions. It is only when all gene modules are surveyed, including those that represent developmental ontogeny, that it is evident how MDM and DAM/ARM-like microglia similarities are superimposed on underlying discrete properties. Our data therefore suggest that a brain macrophage phenotype characterised by a phagolyosomal and lipometabolic transcriptional identity (functional relevance supported as discussed above), and thus resembling chronic disease DAM/ARM, arise after stroke via dual sources i.e. from resident microglia and immigrant monocyte precursors. It is important to emphasise that this phenotype, irrespective of originating cell class, remains distinct from perivascular macrophages, which on global comparison share more similarities with non-reactive microglial clusters (**Figure 7-8**). A recent study showed that brain macrophages expressing MMP12 and osteopontin (encoded by *Spp1*), and thus proposed to reflect key features of the DAM/ARM-like phenotype, were derived from both resident microglia and *Cxcr4*-expressing bone marrow cells^30^. The contribution of dual monocyte and resident brain sources of macrophages accumulating in chronic disease around amyloid deposits has also recently been proposed^77^. By pairwise comparison of differential gene expression between our MDM (cNMG2) and DAM-like (cMG7) microglia, we identified multiplex gene combinations that could be used as *in situ* markers. *Fcrls*^-^*Csf1r*^lo^*Gpnmb*^+^ cells with a highly amoeboid morphology (reflecting *Gpnmb*-expressing MDM) were almost entirely located in the peri-infarct zone which was concordant with the almost exclusive origin of Ly6C^+^ monocyte-derived cells from the MCAO hemisphere in our scRNASeq dataset (Fig 3). These *Gpnmb*^+^ MDM were highly intermixed with peri-infarct *Fcrls*^hi^ microglia and co-located within the same territory as *Fcrls*^hi^*Spp*1^+^ microglia. It reinforces that spatial signals enriched in the peri-infarct zone may shape macrophages in this area towards a conserved phenotype whether of resident brain microglial or blood monocytic origin. This provokes the question as to whether peri-infarct co-located stroke-induced macrophages of distinct ontogenies yet similar effector phenotypes are engaged in similar tasks functionally. Due to the restricted tools available, previous studies have been limited to comparing functional properties of bulk resident microglial and non-resident monocyte-derived populations, but nonetheless have suggested the monocyte-derived cells are more phagocytic^29^. Future studies harnessing advanced tools will be needed to examine comparative functions at more discrete cell state resolution and of functional properties beyond phagocytosis. Moreover, the proximity of microglial- and monocyte-derived macrophages highlights their potential for communication and functional influences on one another. Notably, we previously showed how MDM can affect microglial function *ex vivo* and the impact of monocytes on long-term microglial phenotype after CNS trauma^23^.

There are some limitations and important unanswered questions arising from the current study. The inclusion of ipsilateral and contralateral hemispheres was a productive approach to discover the global brain scRNAseq profile not available to date but can cause challenges in interpreting stroke-specific effects in some cases. However, this is offset by our validation of scRNAseq patterns by *in situ* tissue approaches including sham-operated controls which both showed marked concordance between scRNAseq and ISH patterns and the stroke-specific induction where this may not have been possible to conclude definitively by scRNAseq alone. We studied a single time point 3 d after MCAO which precludes assessment of how the composition of cell states and types may progress into the chronic phase of stroke important for longer-term outcomes. Although pseudotemporal trajectory analyses revealed clear transitional phenotypes it is unclear if some cell states represent transient or discrete fixed phenotypes over a more protracted timeframe and the long-term fate of cells and sub-states remains unknown currently. A recent study of note suggests the DAM/ARM-like microglial state is present chronically after CNS trauma^78^ but whether this results from early conversion then persistence or later but transient conversion is unknown, a distinction that may have considerable implications for therapeutic manipulation. The long-term fate of immigrant sub-populations is also not clear after stroke. We used only young adult male mice and given sex- and age-related differences in myeloid cell properties are increasingly recognised and stroke is more prevalent in older individuals, these factors will need systematically integrated in futures studies. Nonetheless, and in keeping with the broad cross-disease and cross-tissue (i.e. that incorporate differences in age and sex) conservation in profiles of certain cell phenotypes described here, our key findings likely apply irrespective of age and sex. Profiling of human ischaemic stroke tissue will be required to determine which features apply across species and inform new interventional strategies.

In summary, our findings advance understanding of the cellular origins, molecular identities, and spatial patterns contributing to microglial/macrophage diversity after acute stroke. Combinatorial influences of resident and immigrant myeloid ontogeny, and brain location relative to primary stroke infarct pathology and its remote connected sites, create patterns of spatially organised microglial and monocyte-derived cell states. We propose these states will influence global brain function and key stroke outcomes such as cognitive and motor recovery. However, further studies achieving more targeted manipulation of specific myeloid states are warranted to empirically define their functional roles.

## Methods

### Experimental stroke model

Procedures were conducted on male 10-12 week old C57BL/6J mice (Charles River Laboratories). Mice were maintained under specific pathogen-free conditions and a standard 12 h light/ dark cycle with unrestricted access to food and water. Mice were housed in individually ventilated cages in groups of up to five mice and were acclimatised for a minimum of one week prior to procedures. All procedures involving live animals were carried out under the authority of a UK Home Office Project Licence in accordance with the ‘Animals (Scientific Procedures) Act 1986’ and Directive 2010/63/EU and were approved by The University of Edinburgh Bioresearch & Veterinary Services Animal Welfare and Ethics Review Body (AWERB). Experimental stroke was induced by permanent distal middle cerebral artery occlusion (dMCAO) under isoflurane anaesthesia (mixed with 0.2 L/min O_2_ and 0.5 L/min N_2_O). Core body temperature was maintained at 37 ± 0.5 °C throughout the procedure with a homeothermic system (Harvard Apparatus, UK). A vertical incision between the left eye and ear was made and the main trunk and bifurcations of the middle cerebral artery exposed via a subtemporal craniectomy. The site was cooled continuously with saline application. The middle cerebral artery was electrocoagulated at the junction of its main trunk and bifurcation and cessation of blood flow confirmed visually prior to cutting through the coagulated area. The temporal muscle was repositioned, the incision sutured and topical local anaesthetic (lidocaine/prilocaine, LMX4) applied. Anaesthesia was discontinued and mice were administered 0.5 ml sterile saline subcutaneously. Mice were returned to home cages on a heating blanket and then transferred to pre-surgery holding areas.

### Cell sorting and flow cytometry

Mice were perfused transcardially with 0.9% saline under terminal isoflurane anaesthesia to remove circulating blood and brain samples containing the infarct/peri-infarct region or the equivalent region in the contralateral hemisphere were dissected. Samples were placed in 1X Hank’s buffered saline solution (HBSS) (without Ca^2+^ and Mg^2+^) (Gibco, Life Technologies) on ice and minced to fine pieces using a scalpel blade and then transferred to a 15 ml Dounce tissue homogeniser and manually dissociated. Samples were centrifuged at 400 g for 5 min at 4 °C with no brake, supernatant was aspirated and pellets resuspended in 35 % Percoll (GE healthcare) in HBSS (without Ca^2+^ & Mg^2+^) and carefully overlaid with 1X HBSS. Samples were centrifuged at 800 g for 45 min at 4 °C with no brake for density separation of cells from myelin. Supernatant and myelin layer were removed, and cells resuspended in 1X HBSS (without Ca^2+^ or Mg^2+^). Cell suspensions were centrifuged at 400 g for 5 min at 4 °C with no brake and resuspended in staining buffer (DPBS (without Ca^2+^ and Mg^2+^) (Gibco, Life Technologies) containing 0.1 % low endotoxin BSA (Sigma). Cells were incubated with anti-CD16/32 (Clone: 93, Biolegend) for 30 min at 4 °C to block non-specific Fc receptor binding and then for 30 min at 4 °C with the following fluorochrome-conjugated antibodies (anti-CD45-PE (clone 30-F11), anti-CD11b-BV711 (clone M1/70), anti-Ly6G-APC (clone 1A8), anti-CD3-APC-Cy7 (clone 17A2), anti-CD19-PE-Cy7 (clone 6D5), Biolegend) mixed with one of three distinct barcoded hashtag antibody-oligonucleotide (HTO) conjugates (TotalSeq-A, Biolegend) for each mouse to enable cell hashing. Cells were washed, centrifuged 400 g for 5 min at 4 °C with no brake, and resuspended in FACS buffer (0.04% BSA in DPBS). A myeloid-enriched cell population (CD3-CD19-Ly6G-CD45^+^) was sorted into FACS buffer using a FACSAria IIu (Becton Dickinson, Oxford, UK) and cell suspensions stored on ice. Unstained and single-stained controls were used to define the sorting and cytometry analysis gating positions. Approximately 30,000 cells per individual brain sample from each mouse (n = 3 mice) were sorted and cells ipsilateral or contralateral to MCAO pooled i.e. a pooled ipsilateral sample and a pooled contralateral sample each comprising HTO-barcoded cells from three bio-replicates (mice) was formed. Cytometry analysis was conducted on the FCS3 data files acquired during the cell sorting procedure using FCS Express 7 (De Novo Software).

### Single cell library preparation and sequencing

Single cell 3’ gene expression (mRNA) and cell surface hashtag-oligonucleotide (HTO) libraries were prepared according to the Chromium Single Cell 3ʹ with Feature Barcoding technology for Cell Surface Protein Reagent Kit v3 protocol (10X Genomics) with modifications for incorporation of TotalSeq™-A HTO antibodies (clones M1/42; 30-F11, BioLegend). Briefly, generation of gel beads in emulsion (GEM) partitioned with single cells, reverse transcription to generate cDNA and HTO-barcoded DNA, and DNA amplification were performed according to manufacturer instructions with the addition of HTO primers during DNA amplification. Amplified cDNA and HTO-derived DNA were separated by size selection, fragmented and libraries constructed. Expected fragment size distributions for cDNA and HTO libraries were confirmed by electrophoresis (Agilent Bioanalyzer). cDNA and HTO libraries were mixed at a ratio of 4:1 for the pooled sample ipsilateral to MCAO and separately for the pooled sample contralateral to MCAO. Ipsilateral and contralateral samples (each containing combined cDNA and HTO libraries) were loaded on separate lanes of an SP flow cell and paired-end sequencing (Read 1: 28 cycles; i7 index read 1: 8 cycles; Read 2: 91 cycles) was conducted on a Novaseq 6000 (Illumina) to achieve a minimum read depth of 20,000 reads per cell for cDNA (gene expression) and 5000 reads per cell for HTO (mouse ID). cDNA and HTO sequencing reads were demultiplexed according to unique i7 indexes.

### Single-cell RNA-Seq quality control and pre-processing

Count matrices were generated for each library using the 10X Genomics CellRanger v3.0.2 ^79^ pipeline. First, raw reads were demultiplexed to produce fastq files. Second, reads were aligned to the mouse reference genome (GRCm38) and quantified. Resultantly, two unique molecule identifiers (UMI) based expression matrices were generated each for the ipsilateral and contralateral conditions which were pre-processed and analysed using the Seurat v3 package in R ^80^. As the samples were prepared and sequenced together, they represented a single batch and hence analysed together as a merged dataset. Subsequently, quality control was conducted for cells and genes, where genes expressed in at least three cells were incorporated in downstream analyses. Cells were initially filtered based on the number of genes (at least 700) expressed and the mitochondrial percent (less than 8%). Subsequent to TPM like log normalisation using a scaling factor of 10^4^, cells with normalised UMI between 2,500 to 4,000 were considered for downstream analyses. To identify multiplets, HTOs ^42^ were quantified for quality controlled cells of each condition using CITE-seq-Count ^81^ (https://github.com/Hoohm/CITE-seq-Count). The hashtag oligos (HTO) for each cell were normalized using a centred log-ratio transformation. Cells were then demultiplexed using the MULTI-Seq approach ^82^ and those assigned as multiplets or empty droplets were filtered out. To remove donor effects, canonical correlation analysis (CCA) was adopted using default parameters from Seurat ^80^. Briefly, the approach identifies similar cells or “anchors” between datasets (in this case donors) based on their similarity in a joint reduced space, CCA components. The differences between these anchors represents the differences between donors and is accordingly used to calculate a weighted correction vector. These vectors are then applied on cells of the original gene expression matrix to remove donor effects.

### Cell clustering

The integrated scRNA-Seq dataset was then scaled and regressed for percent mitochondrial content. Principle component analysis (PCA) was used to reduce the dimensions of the dataset to the 47 most significant (*P* < 0.05) PCs based on Jackstraw permutations ^83^. Clustering on the whole dataset was done using a network-based approach using Graphia^84^. A k-nearest neighbour (k = 10) network of cells was constructed by connecting cells with a high Pearson similarity coefficient (based on the significant PCs) r ≥ 0.5 to accommodate all cells. The resulting network was clustered using the Markov Clustering algorithm (MCL) with a low inflation MCLi = 1.35 ^85^.

### Comparing cell proportions

The scRNA-Seq experiment was conducted by enriching for myeloid cell types, and the considerably altered composition of mononuclear phagocytes in response to MCAO made a direct comparison between MCAO and CTRL relative abundance inaccurate. Hence, we assumed that the number of resident homeostatic or homeostatic-like microglia from cluster 1 and 2 respectively would not change significantly between conditions, therefore we normalised cell frequencies of each cluster (considering condition and donor) with those of cluster 1 and 2. The normalized cell frequencies were then compared between conditions for cell clusters using a t-test.

### Generating pseudo-bulk samples from scRNA-Seq to construct gene co-expression networks

All the gene co-expression networks were constructed by first constructing pseudo-bulk samples from the scRNA-Seq data and then conducting weighted gene co-expression network analysis or WGCNA ^86^. This preceding step of constructing pseudo-bulk samples aimed to improve the signal-to-noise ratio of single-cell data, capture the intra- and inter-cell type variation and generate a cell type balanced correlation space i.e. each cell type is equally represented as each has equal number of representative pseudo-bulk samples. The pseudo-bulk algorithm was used to generate five pseudo-bulk samples for each cluster. The strategy for generating these pseudo-bulk samples was conducted in the following two steps. 1) Each of the clusters was further clustered into five sub-clusters using Clara^87^ (an extension of k-medoid clustering) based on the cell-to-cell similarity determined by the significant PCs. In the process, the medoid cell of each sub-cluster was also identified. 2) For each of these representative medoid cells, the expression values for genes were averaged across their ten nearest neighbours (based on significant PCs) within the cluster. As a result, five sub-clusters/pseudo-bulk samples were generated for each of the original clusters. The resultant pseudo-bulk vs gene expression matrix was used to identify co-expressing genes using WGCNA ^86^. A soft threshold of = 6 was used to construct an adjacency matrix of pairwise Pearson correlations between genes. This matrix was then transformed into a topological overlap matrix to take into account the structure of the data. Hierarchical clustering was performed with parameters set for the module size (minModuleSize = 10), the granularity of clustering (deepSplit = 4) and the number of clusters (mergeCutHeight = 0.15).

### Functional enrichment analysis of gene clusters

To provide functional annotation to gene clusters generated from WGCNA, the clusters were examined for their enrichment in biological processes as defined by gene ontology (GO)^88^, Reactome^89^ and Kyoto encyclopedia of genes and genomes (KEGG)^90^ databases using clusterProfiler in R ^91^. Furthermore, to compare the enrichment of KEGG terms across gene clusters, the top 6 terms for each cluster were compared across all clusters shown in Figures. In all cases, the biological processes with significant (*Padj* < 0.05) enrichment were considered with *P* adjusted using the Benjamini-Hochberg correction for multiple testing.

### Cell cycle analysis

Cell clusters were annotated for one of the three cell cycle stages, including G1, G2 and S phase using the cell cycle scoring algorithm available in Seurat v3 ^80^ adapted from ^45^. The algorithm generates a standardised score for each cell based on their average expression of G2 and S phase gene signatures. Based on these scores, cells are assigned to one of the three phases, where G1 (non-cycling) phase presents poor scores for both G2 and S phase gene signatures.

### RNA velocity and pseudotime analysis

To estimate the direction and velocity of differentiating cells, the velocyto package in python was used with parameters proposed by the authors (https://github.com/velocyto-team/velocyto-notebooks/blob/master/python/Haber_et_al.ipynb). First, the spliced and unspliced reads were quantified from the BAM files generated from the cellranger pipeline. Cells previously selected based on the Seurat QC were used in the analysis. Genes were filtered based on their minimum expression, average expression and coefficient of variation. The data was subsequently normalized, and the 3,000 most variable genes were used to calculate PCAs. The top nine PCs were then used for kNN-imputation with k = 70 and a gamma distribution was fitted to each gene. The resultant expression matrix was used to estimate the velocity and future cellular states i.e. the direction.

Highly variable genes along the differentiation trajectory were determined using pseudotime analysis using Slingshot with default parameters ^59^. First, the algorithm re-clustered the data using a gaussian mixture model. Next, the start and end points of differentiation previously determined by RNA velocity were used to construct a minimum spanning tree on which simultaneous principal curves were fitted. Each cell was then aligned to the curves, thus aligning them along pseudotime. Finally, genes significantly changing (*P* < 0.05) along pseudotime were determined using a general additive model.

### Cross-model microglial comparison

Nine studies (including this study) were selected to represent different neurological disease conditions for the cross-model microglial comparison ^46-53^. Microglial clusters from within these studies were identified and included for the analysis. As a note, DEG analysis for most studies compared one microglial cluster to all others, while DEGs from Mathys et al. ^47^ were identified by comparing early/late response microglial subtypes (Cluster 3, 6 and 7) with homeostatic microglia (Cluster 2). To capture genes representative of a cluster, DEGs were filtered by log fold change > 0.25, and Padj < 0.05. In the case of Mathys et al. ^47^, genes with maximum likelihood estimation > 0 (alternate metric to fold change) were considered. Furthermore, of these filtered DEGs the top 500 genes (based on log fold change) were considered for each microglial cluster across publications. The resultant gene lists were compared using Fischer’s exact t-test which provided an odds ratio (values greater than 1 are indicative of associated/correlated gene lists) and a *P*. Furthermore, only significant (P < 0.05 and odds ratio >1) pairwise odds ratios were examined.

### Differential gene expression

Differentially expressed genes for each cluster were identified using the negative binomial generalized linear model available in Seurat^92^. Furthermore, only those genes were tested whose log fold change was greater than 0.25 (in either direction) and expressed in a minimum of 25% of cells in the groups examined.

### Multiplex fluorescence *in situ* hybridisation

RNAScope™ Multiplex Fluorescent Assay v2 (ACDBio, 323100) was performed on 20µm cryostat-sectioned fixed-frozen brain sections using slight modifications to manufacturer’s protocol. Sections were allowed to equilibrate to room temperature (RT) before washing with dH_2_O and baking on to slides at 60°C. Slides were post-fixed (10% neutral-buffered formalin) for 15 min at 4°C and dehydrated through alcohols. Hydrogen peroxide was applied for 10 min at RT and antigen retrieval was performed for 15 min in a pre-heated (20 min) plastic Coplin jar in a 97.5°C waterbath. Sections were left at RT overnight before placing into humidity chambers and protease III was applied for 30 min at 37°C. The following probes were incubated for 2 hours at 40°C *Fcrls* (441231-C2)*, Ccl3* (319471)*, Spp1* (435191-C3)*, Csf1r* (428191)*, Gpnmb* (489511-C3)*, Ccr2* (501681)*, Mki67* (416771-C3). Probes were subsequently amplified and labelled with Tyramide 488 (Invitrogen, B40953), 555 (Invitrogen, B40955) or 647 (Invitrogen, B40958). Nuclei were labelled with 1µg/mL DAPI. Low-power images were acquired using an Axioscan Slide Scanner and high-power confocal images using an Opera Phenix Plus high-content confocal imager. Maximum intensity projections were processed from confocal stacks and cells from 3 – 6 non-overlapping 300µm x 300µm regions of interest were manually quantified per brain region per section.

### Immunofluorescence

For immunolabelling of P2Y12 and Ki-67, sections were equilibrated to RT, washed with dH_2_O and baked for 30 min at 60°C. Antigen retrieval was performed using Tris-EDTA buffer (10mM Tris Base, 1 mM EDTA, pH 8.6) for 20 min in a non-preheated plastic Coplin jar in a water bath set to 97.5°C for 20 min. Rabbit-anti-Ki-67 (1.25µg/mL, ab15580) was incubated overnight at 4°C. Endogenous peroxide activity quenched with 0.3% H_2_O_2_ RT 30 min. Biotinylated Goat-anti-Rabbit (ZH0615) incubated for 90 min, then strep-HRP (Tyramide amplification kit, B40933) incubated for 45 min before amplification with Tyramide 555. To allow multiplexing of antibodies raised in the same species, antigen retrieval was performed again in the same manner to elute Ki-67 antibodies (leaving Tyramide signal untouched). Rabbit-anti-P2Y12 (AS55043A) was incubated overnight at 4°C and fluorescently labelled using Goat-anti-Rabbit-AlexaFluor-647 (A21244), incubated for 90 min RT. Primary antibodies were incubated in buffer containing 1% BSA, 0.3% Triton-X 100, 1X PBS, other antibodies were incubated in 0.1% BSA, 0.5% Tween, Tris-buffered-Saline. 3 x 2 min washes were performed between each step following primary antibody incubation using 0.5% Tween, Tris-buffered-Saline. Widefield images of P2Y12 and Ki67 were acquired using an AxioImager.D2.

### Statistical analysis

Our primary aim was to conduct a hypothesis-free scRNASeq profiling study. Formal sample size estimates were not conducted however cell isolation was guided by estimating the number of cells required to detect a rare cell type within each individual sample using the following parameters (frequency = 0.01, minimum number of cells = 10, detection power = 0.95) (https://satijalab.org/howmanycells/)93. Cell hashing (see above) from three independent mouse brains provided bio-replication within the dataset. Statistical approaches for analysis of scRNA data are described above. Cytometry data were analysed by Welch’s *t*-test using Graphpad Prism (v9). For quantification of multiplex smFISH, a linear mixed-effect model was utilised to compare the distribution of cell phenotypes within brain regions across sham and stroke brains. Within-subject design with random intercepts was used and factors (phenotype, region) and interactions were treated as fixed-effects and their inclusion evaluated through log-likelihood ratio. Holm-Sidak post-hoc tests were utilised for a-priori planned pairwise contrasts. Normality and homoscedasticity were evaluated graphically and box-cox transformations applied when necessary. One sided t-tests were utilised to compare stroke phenotypes to 0 counts seen in sham controls. A paired t-test was used to compare phenotype abundance within stroke brains and all p values were adjusted using the Bonferroni method. All smFISH analyses were performed using R (Version 4.1.2).

## Data and code availability

Data and code are available upon reasonable request to the authors.

## Acknowledgments

We acknowledge funding from the Medical Research Council (MR/R001316/1, MR/L003384/1) (AA, BWM), the Leducq Foundation Transatlantic Network of Excellence, Stroke-IMPaCT (19CVD01) (BWM), Alzheimer’s Research UK (ARUK-PG2016B-6) (KH, BWM), the UK Dementia Research Institute which receives its funding from the Medical Research Council, Alzheimer’s Society, and Alzheimer’s Research UK (SS, LL, BWM), Alzheimer’s Research UK Scotland Network (NCH, KH, BWM), Chief Scientist Office Scotland (CGA/18/46) (AO-A, BWM), Stroke Association (TSA PPA 2017/01) (JB). NCH is supported by a Wellcome Trust Senior Research Fellowship in Clinical Science (ref. 219542/Z/19/Z). We thank staff at Edinburgh Genomics, FACS and biological research facilities for their support to the studies. For the purpose of open access, the author has applied a Creative Commons Attribution (CC BY) licence to any Author Accepted Manuscript version arising from this submission.

## Author contributions

Conceptualization: PR, BWM; Methodology: all authors; Investigation: all authors; Analysis: all authors; Supervision: BWM; Project administration: BWM; Writing – original draft: AP, JB, BWM; Writing – review and editing: all authors; Funding acquisition: NCH, KH, BWM.

## Supplementary information

### Supplementary Figures

Figure S1. Flow cytometry analysis gating Figure S2. Cell sorting gating

Figure S3. Multiplet removal from scRNAseq data using HTO

Figure S4. Top 5 differentially expressed genes for all cell clusters

Figure S5. Top 5 differentially expressed genes for sub-clustered microglia

Figure S6. Gene coexpression network analysis of microglial states

Figure S7. Top 5 differentially expressed genes for non-microglial mononuclear myeloid clusters

Figure S8. Gene coexpression network analysis of non-microglial mononuclear myeloid cells

Figure S9. Gene coexpression network analysis of all mononuclear myeloid cell clusters

Figure S10. Proteogenomic profiles of MCAO-associated myeloid cells

Figure S11. Spatial mapping of reactive microglia in sham controls

### Supplementary Tables

Table S1. Overall differential gene expression analysis

Table S2. Overall cluster annotations and top differentially expressed genes

Table S3. Subclustered microglia annotations and differentially expressed genes

Table S4. Subclustered microglia differential gene expression analysis

Table S5. Subclustered microglia WGCNA gene coexpression clusters

Table S6. Subclustered microglia enrichment analysis of gene coexpression clusters

Table S7. Studies used in cross-model comparison of microglia

Table S8. Subclustered non-microglia, differential gene expression analysis

Table S9. Subclustered non-microglia WGCNA gene coexpression clusters

Table S10. Subclustered non-microglia cluster annotations & genes

Table S11. Subclustered non-microglia enrichment analysis of gene coexpression clusters

Table S12. Integrated subclustered myeloid differential gene expression analysis

Table S13. Integrated subclustered myeloid WGCNA gene coexpression clusters

Table S14. Integrated subclustered myeloid enrichment analysis of gene coexpression clusters

Table S15. Differential expression cMG7 vs cNMG2 from integrated subclustered myeloid analysis

