## Supplementary Figures for "Phenotypic and spatial heterogeneity of brain myeloid cells after stroke is associated with cell ontogeny, tissue damage, and brain connectivity"

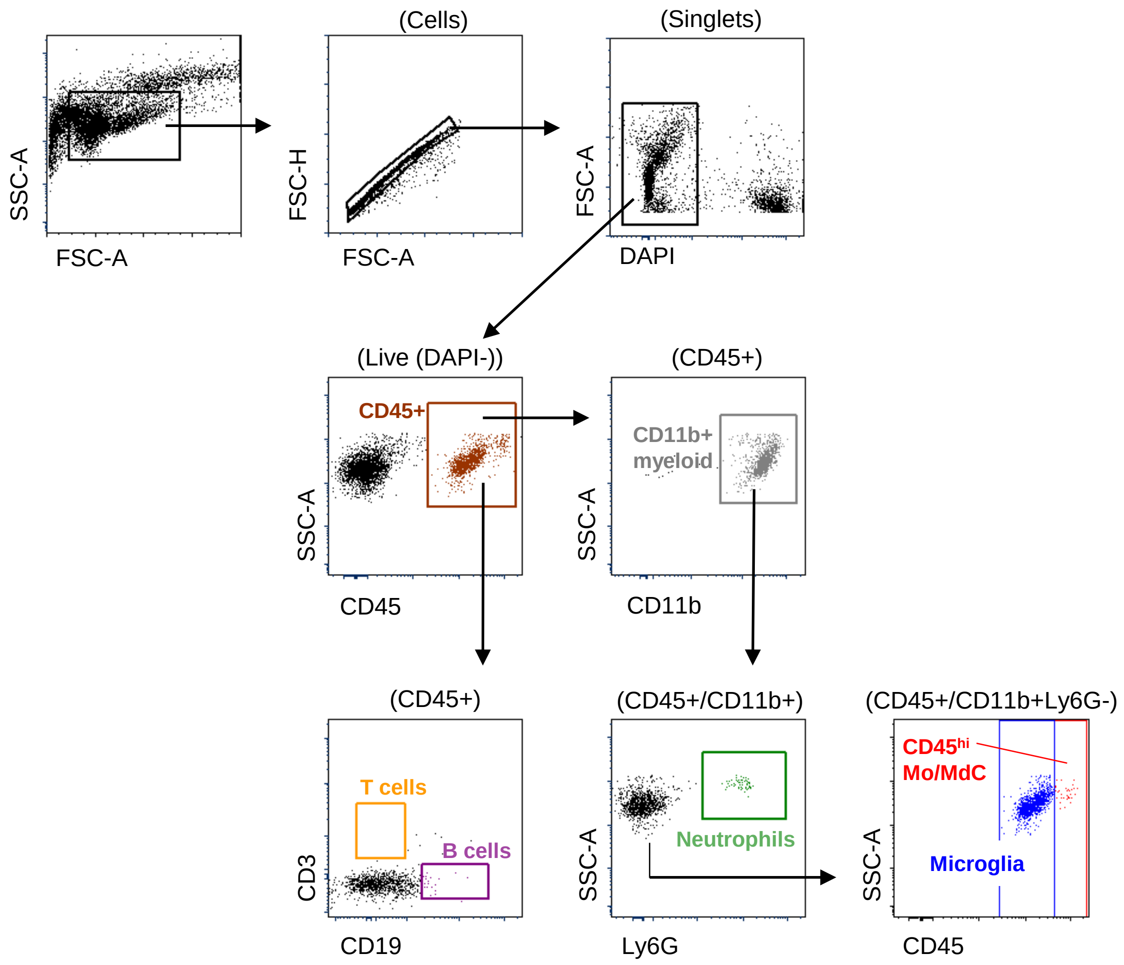


**Figure S1. Flow cytometry analysis gating**

Gating scheme applied to brain cell suspensions prepared from ipsilateral (MCAO) and contralateral (CTRL) hemispheres 3 d after MCAO to identify major immune cell populations. $CD45^{+}$ cells were identified and gated for major lymphocyte populations using CD3 (T cells) and CD19 (B cells) expression, and major myeloid cell populations using combinations of CD11b, Ly6G and CD45 expression to define neutrophils ($CD11b^{+}Ly6G^{+}$), microglia ($CD11b^{+}Ly6G^{-}CD45^{lo}$), and monocytes (Mo)/monocyte-derived cells (MdC) ($CD11b^{+}Ly6G^{-}CD45^{hi}$).

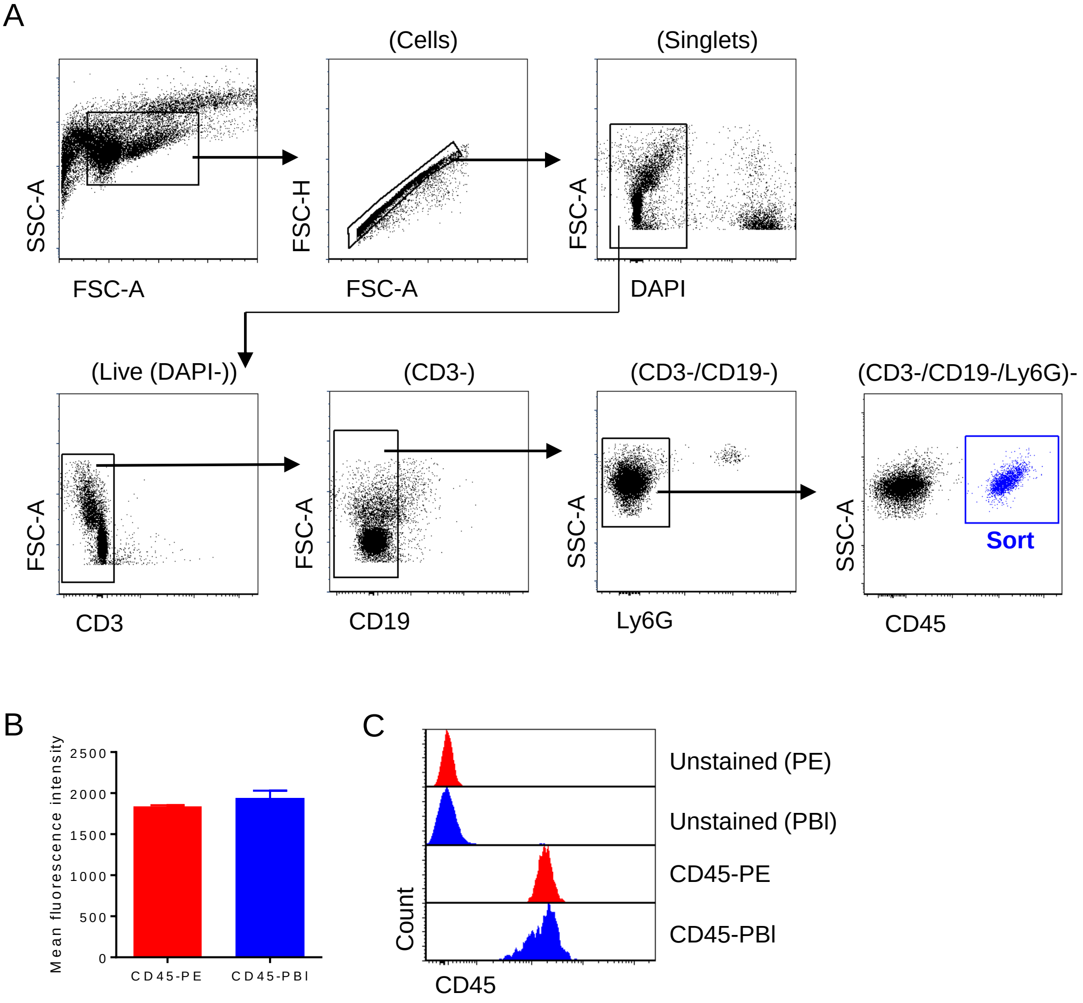


**Figure S2. Cell sorting gating**

(A) Gating scheme applied to brain cell suspensions prepared from ipsilateral (MCAO) and contralateral (CTRL) hemispheres 3 d after MCAO to isolate a mononuclear myeloid-enriched cell population for scRNAseq. Cells negative for CD3, CD19 and Ly6G, and positive for CD45 were sorted to exclude the majority of T cells, B cells and neutrophils. (B) Mean fluorescence intensity and (C) representative histograms of CD45 intensity for cells dual labelled with CD45 antibodies conjugated to different fluorochromes.

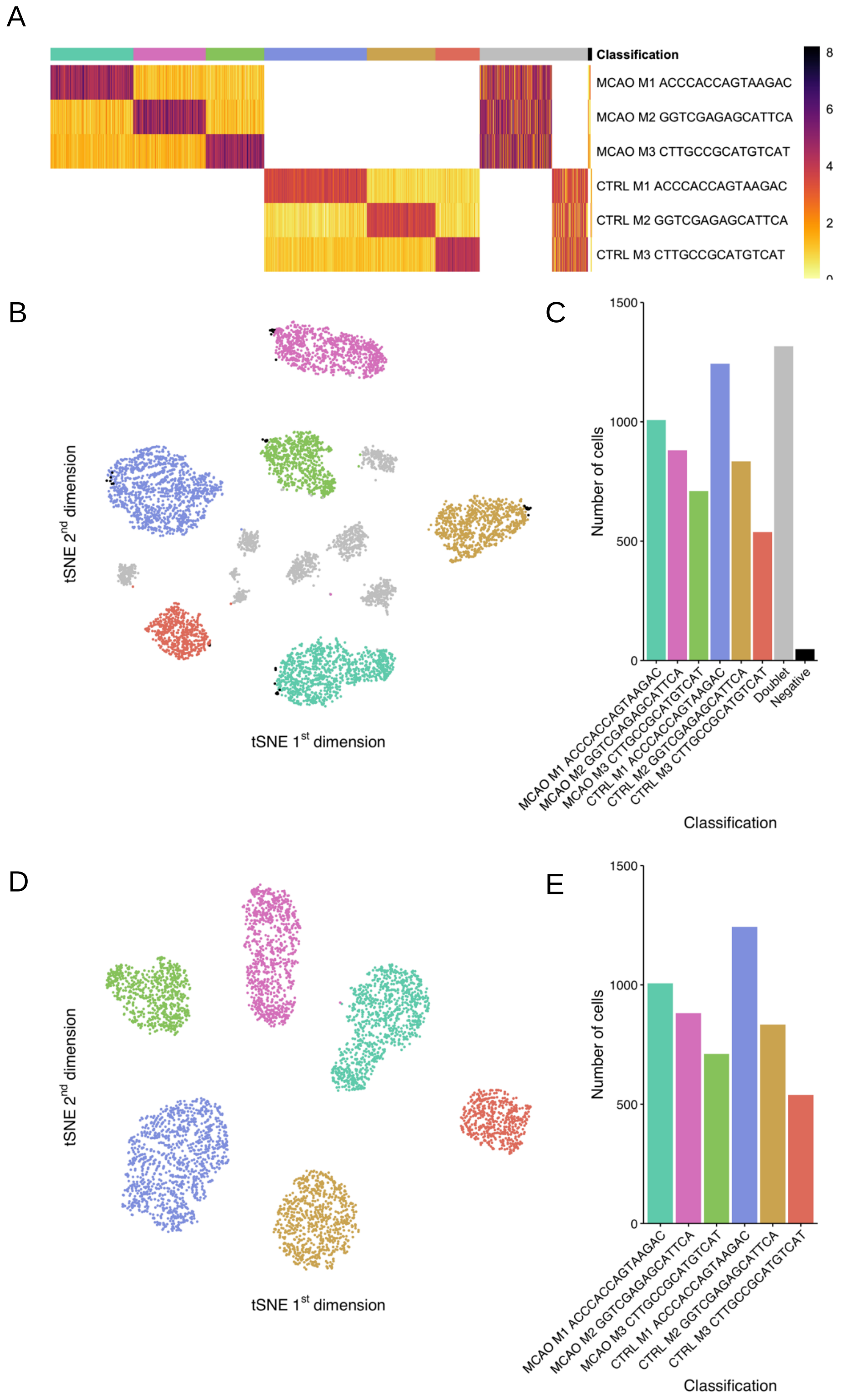


**Figure S3. Multiplet removal from scRNAseq data using HTO**

(A) Heatmap of cells and corresponding hashtag oligonucleotide (HTO) levels for each of the three donors (M1, M2 and M3) across the MCAO and CTRL samples, respectively. Multi-Seq demultiplexing of cells based on donor specific HTOs. Cells are assigned to either a specific donor, multiple donors (multiplets) or none of the donors. The heatmap is ordered based on the assignment of cells (n = 6,576). (B) Cells are projected into 2D space using UMAP based on their normalized levels of HTO. Colours of cells represent their donor classification. (C) Frequency of each classification is shown as a bar plot. On removing multiplets and negatively classified cells in a similar fashion the (D) cell UMAP and (E) frequency bar plot is shown.

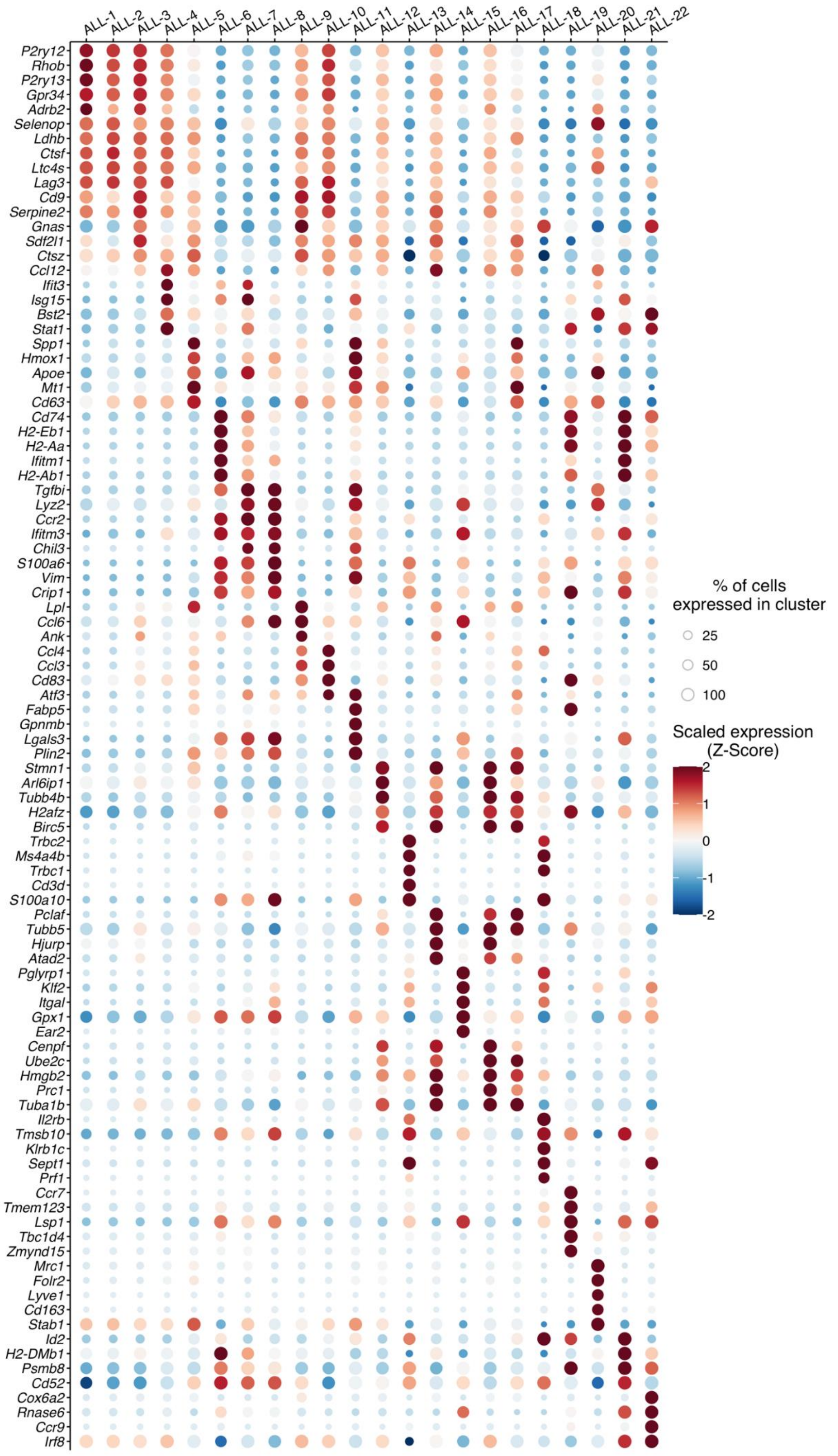


**Figure S4. Top 5 differentially expressed genes for all cell clusters**

Dot plot showing expression of the top 5 differentially expressed genes for each cell cluster (column) (log FC > 0, P< 0.05 versus all other clusters). The colour of each dot represents the scaled average expression and the size of the dot the proportion of cells expressing the gene within the cluster.

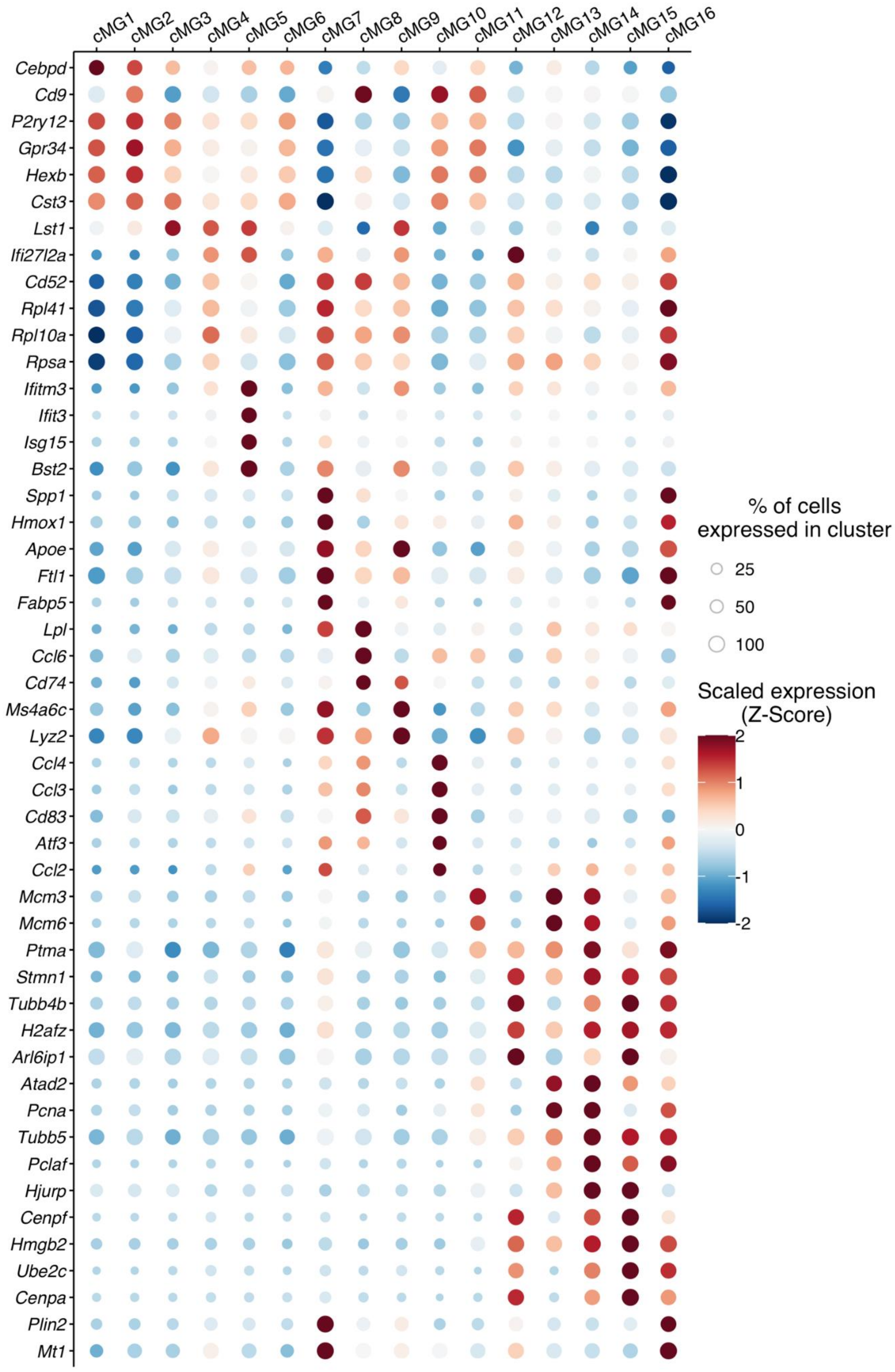


**Figure S5. Top 5 differentially expressed genes for sub-clustered microglia**

Dot plot showing expression of the top 5 differentially expressed genes for each microglial cell cluster (column) (log FC>0, P < 0.05 versus all other sub-clustered microglial clusters). The colour of each dot represents the scaled average expression and the size of the dot the proportion of cells expressing the gene within the cluster.

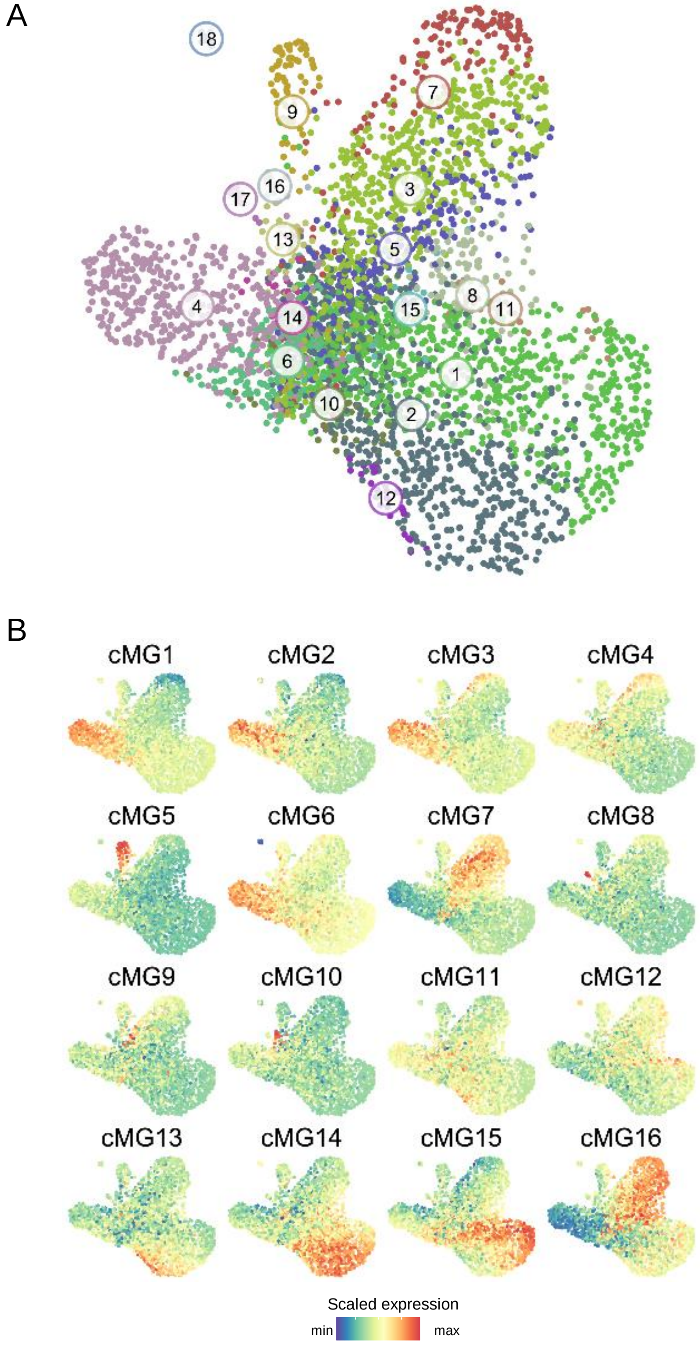


**Figure S6. Gene coexpression network analysis of microglial states**

(A) Gene coexpression network comprising all genes differentially expressed across microglial clusters. Each dot represents a gene and proximity to other genes reflects similarity in transcriptional profile across all sub-clustered microglial cells. Coexpressed genes forming distinct gene modules identified using WGCNA share the same colour with each gene module numbered as shown. (B) Each panel shows the expression level (scaled) of the genes (and gene modules) comprising the coexpression network applied to each microglial cell cluster (**Figure 4A**). Highly expressed genes are in red while lowly expressed genes are in blue. The plots demonstrate how each microglia cell cluster is formed through differential and modular gene coexpression (NB the heatmap in **Figure 4D** summarises these data to show the average expression level for all coexpressed genes within each gene module across cell clusters).

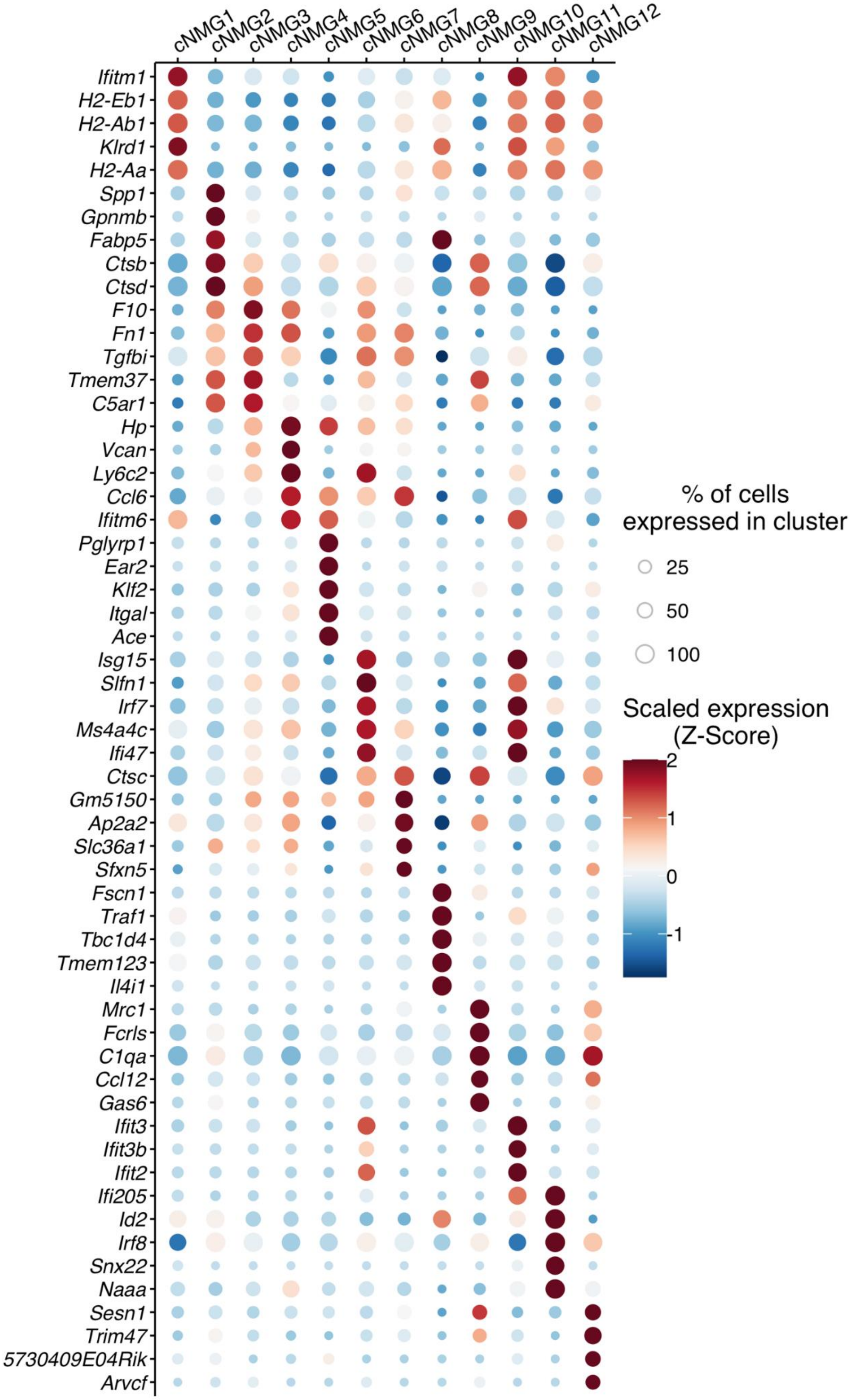


**Figure S7. Top 5 differentially expressed genes for non-microglial mononuclear myeloid clusters**

Dot plot showing the expression of the top 5 differentially expressed genes for each non-microglial mononuclear myeloid cell cluster (column) (log FC > 0, P< 0.05 versus all other sub-clusters). The colour of each dot represents the scaled average expression and the size of the dot the proportion of cells expressing the gene within the cluster.

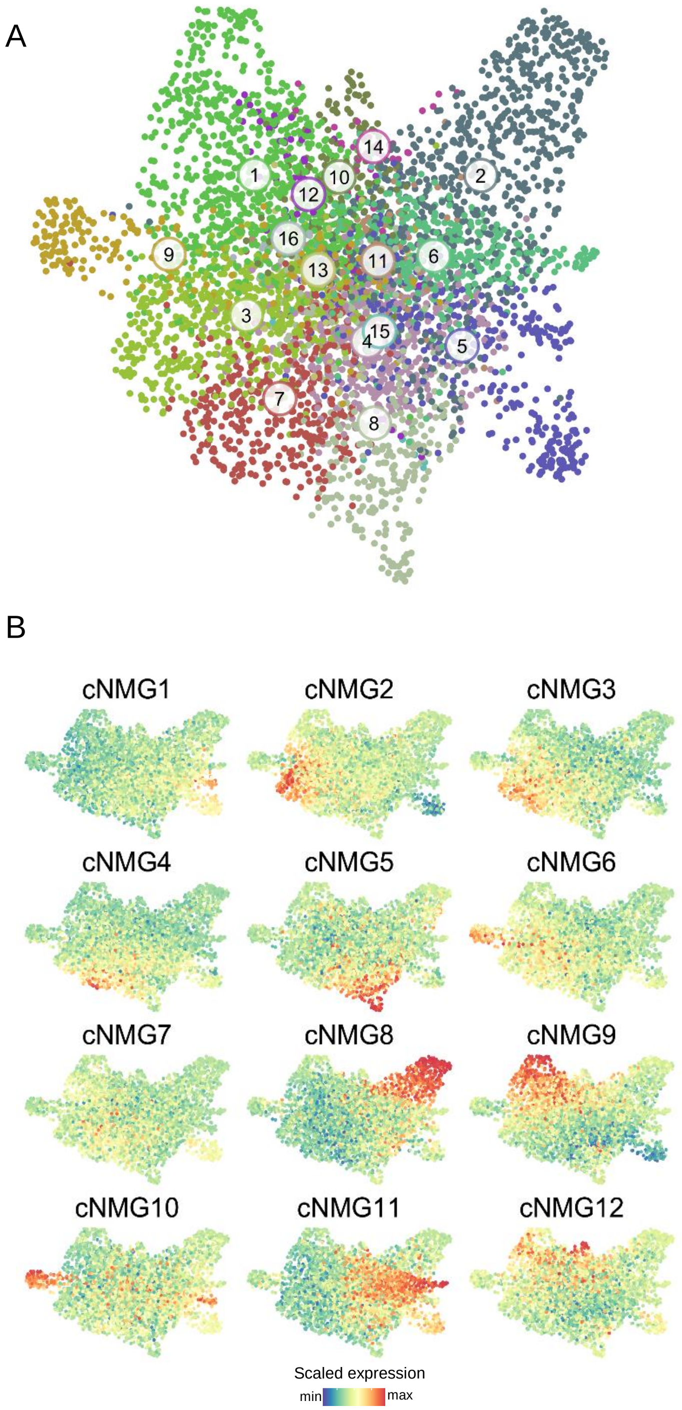


**Figure S8. Gene coexpression network analysis of non-microglial mononuclear myeloid cells**

(A) Gene coexpression network comprising all genes differentially expressed across non-microglial myeloid clusters. Each dot represents a gene and proximity to other genes reflects similarity in transcriptional profile across all cells. Coexpressed genes forming distinct gene modules identified using WGCNA share the same colour with each gene module numbered as shown. (B) Each panel shows the expression level (scaled) of the genes (and gene modules) comprising the coexpression network applied to each non-microglial myeloid cell cluster (F**igure 7A**). Highly expressed genes are in red while lowly expressed genes are in blue. The plots demonstrate how each cell cluster is formed through differential and modular gene coexpression (NB the heatmap in **Figure 7D** summarises these data to show the average expression level for all coexpressed genes within each gene module across cell clusters).

A

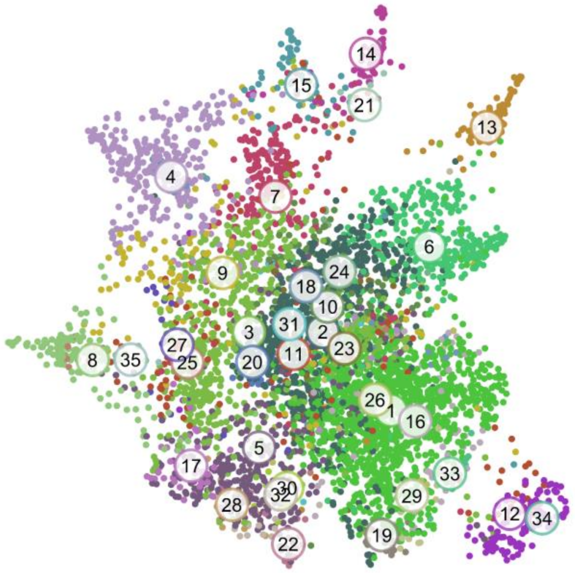


B

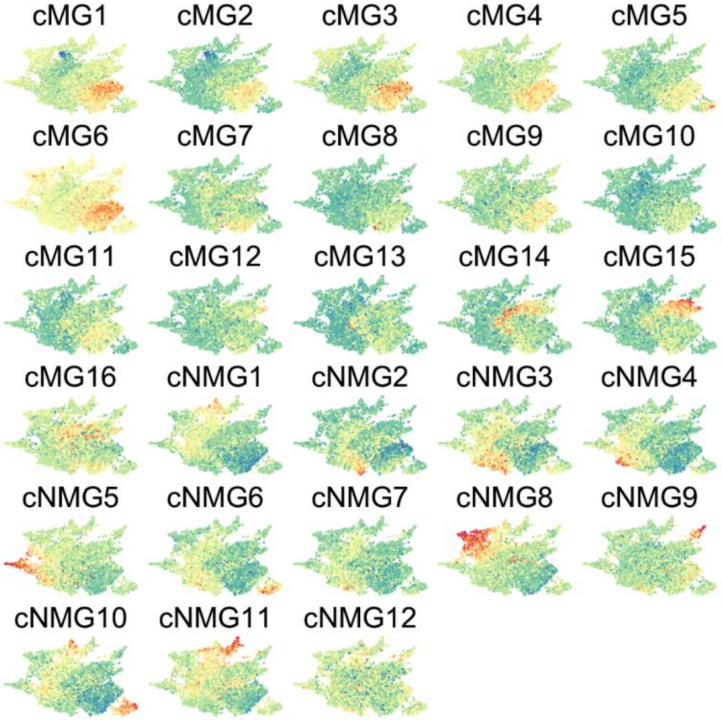


**Figure S9. Gene coexpression network analysis of all mononuclear myeloid cell clusters**

A) Gene coexpression network comprising all genes differentially expressed across the composite of microglial and non-microglial myeloid clusters (see **Figure 8A**) obtained by sub-clustering the original dataset. Each dot represents a gene and proximity to other genes reflects similarity in transcriptional profile across all cells. Coexpressed genes forming distinct gene modules identified using WGCNA share the same colour with each gene module numbered as shown. (B) Each panel shows the expression level (scaled) of the genes (and gene modules) comprising the coexpression network applied to cell cluster (see **Figure 8A**). Highly expressed genes are in red while lowly expressed genes are in blue. The plots demonstrate how each cell cluster is formed through differential and modular gene coexpression (NB the heatmap in **Figure 8B** summarises these data to show the average expression level for all coexpressed genes within each gene module across cell clusters).

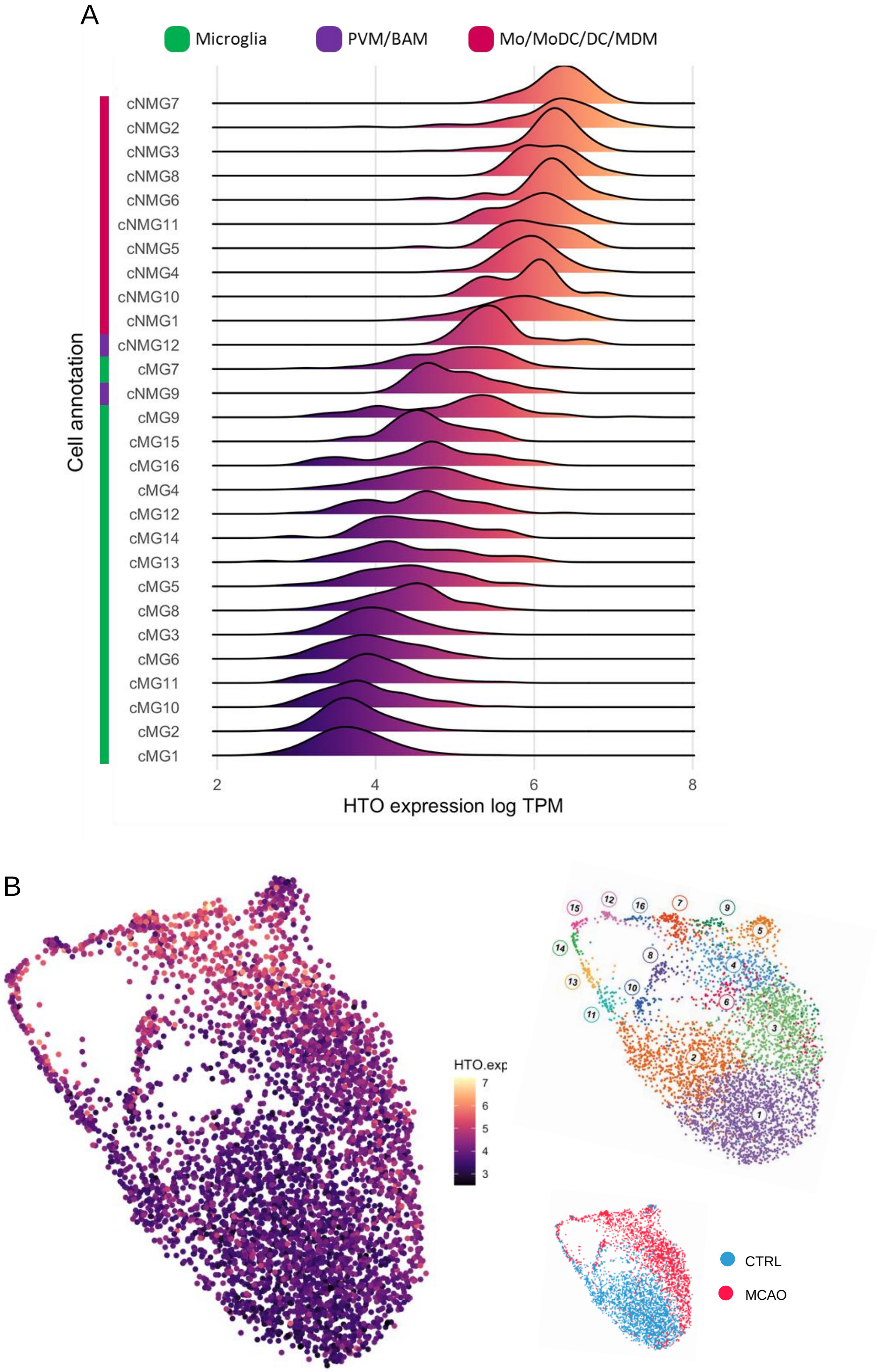


**Figure S10. Proteogenomic profiles of MCAO-associated myeloid cells**

(A) Hash-tag oligonucleotide-derived expression (HTOseq) intensity reporting cell surface CD45 protein levels within each mRNAseq-defined cell cluster from **Figure 8A**. Cell clusters are ordered top to bottom according to median HTOseq expression level (high to low). (B) UMAP of sub-clustered microglia (from **Figure 4A**) showing HTOseq expression levels on each cell. Insets on right show mRNAseq-derived cell clustering (top) and sample of origin (bottom) for reference. PVM, perivascular macrophage; BAM, border-associated macrophage; Mo, monocyte, MoDC, monocyte-derived dendritic cell; DC, dendritic cell; MDM, monocyte-derived macrophage.

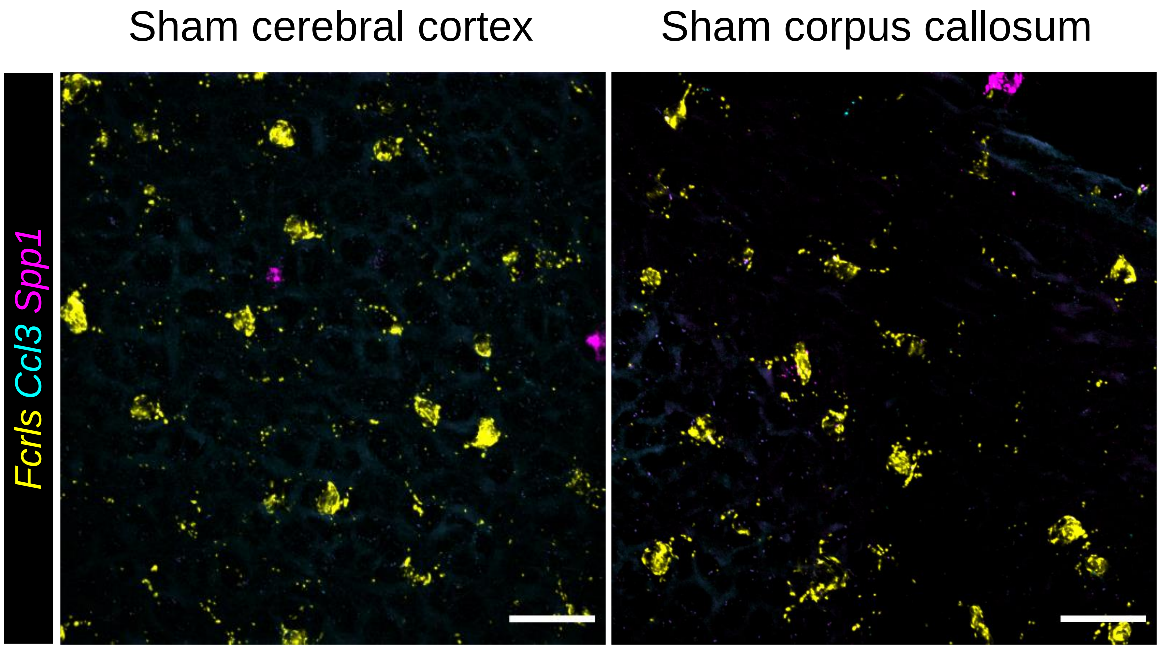


**Figure S11. Spatial mapping of reactive microglia in sham controls**

Multiplex fluorescence *in-situ* hybridisation for *Fcrls*, *Ccl3* and *Spp1* in (left) cerebral cortex and (right) corpus callosum (CC) of Sham-operated control mouse brains. One representative image of four biological replicates. Scale bar = 50µm.
